# IRF7 impacts on prostate cancer cell survival in response to radiation

**DOI:** 10.1101/2022.09.23.509205

**Authors:** Adam Pickard, Francesca Amoroso, Kelsey McCulloch, Andrew Erickson, Ashwin Sachdeva, Rebecca Steele, Debayan Mukherjee, Margaret Dellett, Jonathan McComb, Laura McCaffery, Claire A. Hart, Michael D. Brown, Simon McDade, David Waugh, Noel Clarke, Karl Butterworth, Tim Illidge, Tuomas Mirtti, Ian M. Overton, Ian G. Mills

## Abstract

Understanding the impact of radiotherapy on the evolution of treatment resistant prostate cancer is critical for selecting effective treatment combinations. Whilst activation of Type 1 interferon signalling is a hallmark of how cells respond to viral infection, in cancer cells, multiple stresses are known to activate this same response. In this study we have evaluated for the first time the changes in the interferon response induced by culturing prostate cancer cells under sphere-forming conditions and following irradiation. We report a conserved upregulated transcript profile for both conditions that is strongly associated with therapeutic resistance and cell survival in vitro and in vivo. The profile includes and is regulated by the Type 1 interferon master regulator IRF7 which, when depleted, delays tumour re-growth following irradiation. We immuno-stained two independent prostate cohorts for IRF7 and found that increased expression, particularly in cases with low PTEN expression, correlated with poor prognosis. To more comprehensively characterise the impact of IRF7 and radiation on cells, RNA-Seq was performed on IRF7 knockdown cells at different radiation doses. We identified a number of biological processes that were IRF7-dependent, including the formation of stem-like cell populations and also therapeutic vulnerabilities. For example, irradiation sensitised surviving cells to either a combination of an IKKε/TBK1 and a MEK inhibitor or treatment with an inhibitor of IDO1, an IRF7- dependent gene. Translationally our work suggests that IRF7 expression can be used to stratify patients who may not benefit from receiving radiotherapy alone but rather may benefit from treatment combinations. In two cohorts treated with radical intent, strong IRF7 staining was associated with disease-specific death implicating this pathway as a convergence point for therapeutic resistance in prostate and potentially other cancer types.

## Introduction

Prostate cancer (PCa) is the most common male cancer with 20-30% of cases progressing to metastatic disease and is a leading cause of cancer death in males worldwide [1]. The main therapy for patients is surgery or radiotherapy and in metastatic disease hormonal therapy that targets the AR [2] [3] [4]. However, therapeutic resistance in metastatic disease often develops within 18 months [5]. As few treatment options are available for hormone refractory PCa, understanding how therapy resistant disease develops is therefore key. In addition, approaches which could identify patients likely to develop metastatic PCa could allow earlier intervention and improve survival rates. The development of resistance is associated with the adaptation of signalling pathways which override the normal mechanism of drug action. PTEN-loss constitutes a subtype of PCa linked to both poor patient outcome and therapy resistance [6] [7], which may also be independent of genotype. In fact, therapeutic challenge itself can result in the selection for multidrug resistant cells [8]. In PCa, stem cells have been demonstrated to be multidrug resistant [9], and whilst they represent less than 0.1% of the tumour epithelium, they are androgen receptor negative [10], and are proposed as a mechanism of resistance to anti-androgen therapies [11]. Cancer stem cells are also often described as tumour-initiating cells [12] with small numbers of cells being able to form tumours in in vivo models [13]. Moreover, the selection of stem-like populations is known to contribute to radiotherapy resistance [14]. Therefore, to understand how tumours recur and develop therapeutic resistance, we need to evaluate how cells can perceive multiple cellular stresses and integrate this into a pro-survival signal. How cancer cells respond to cellular damage following treatment may involve development of new adaptive signalling pathways but may also rely upon pre-established biological processes and pathways. In response to viral infection activation of the Type-1 interferon signalling is observed [15], induced by the recognition of viral DNA and RNA mediated by cytoplasmic receptors, such as RIG-I (DDX58), MDA5 and protein kinase R (PKR). These commonly lead to the activation of the interferon stimulatory factors, IRF3 and/or IRF7, and NF-κB and induction of interferon alpha and beta expression [16]. In this study we associate this with cells that survive radiotherapy and are responsive to IKK/MEK and IDO inhibitors. We investigate the association of IRF7 expression with PTEN expression in clinical samples and identify a poor-prognosis subset defined by high nuclear IRF7 expression and low PTEN expression. This work has implications for patient stratification and prediction of treatment response.

## Results

In order to understand how cancer cells may acquire survival phenotypes and develop therapeutic resistance, we interrogated publicly available datasets and analyzed the gene expression of irradiated surviving cells or those that were able to form spheroids in low mitogen growth condition [13] [17]. We initially identified 81 genes that were overexpressed in PC3, DU145 and HCT116 cells grown under sphere-forming conditions versus in monolayer cultures (**Figure 1A**). Given that among all the prostate cancer cell lines available, DU145 and PC3 cells are notoriously the most resistant to radiation, we went on to identify genes that were overexpressed in PC3 cells that survived fractionated regimes (10×1Gy or 5×2Gy) compared to single high-dose exposures (10Gy). We identified 66 genes that were overexpressed following fractionated exposures but not following single-dose exposure (**Figure 1B**). Fourteen genes were overexpressed in all of the sphere-forming conditions and the low-dose fractionated conditions - DDX58, DDX60, IFIT1, IFIT3, IFI44, HERC6, ISG15, OAS1, OAS2, IFI6, IFIH1, PARP9, IRF9 and IRF7 (**Figure 1C**). The 14 upregulated transcripts which included the transcription factors IRF7 and IRF9, were independently validated in PC3 cells by real-time PCR in both spherogenic conditions and upon treatment with fractionated doses of radiation (3×8Gy and 5×2Gy) (**Figure 1D- E**, **Supplemental Figure 1A**). Gene Ontology analysis highlighted a role of cytokines and Type-I interferon signaling when we considered both cells grown in sphere-forming conditions versus in monolayer cultures (**Supplemental Figure 1B**) and PC3 cells that survived fractionated regimes (10×1Gy or 5×2Gy) compared to single high-dose exposures (10Gy) (**Supplemental Figure 1C**). The Type-I interferon pathway is normally activated in response to viral infections, however other stresses, including spherogenesis, irradiation and ATM mutations are also known to activate this pathway [13] [18] [19] [20] [21]. Interestingly, with increasing radiation doses and cell growth in spheroid conditions, we also observed in PCa cell lines a significant increase in the proportion of cells expressing putative stem-like markers (CD133 and CD44) by 60% to 90% respectively (**Figure 1F-G** and **Supplemental Figure 1D**). To determine whether the overexpression of these 14 genes and the expression of stem-like markers is a conserved biology, we irradiated a panel of PCa cell lines (**Supplemental Figure 1E**). We observed a strong linear correlation between irradiation, the expression of these 14 genes and stem-like markers in LNCaP, 22Rv1 and PC3 cells (p<0.05), implying that this is a conserved biology that is activated when the PCa cells undergo stress. The strong association of the 14 transcripts and the stem-like population led us to hypothesize that these genes may regulate these phenotypes. The strongest amongst these 14 genes was IRF7, which is an established master regulator of the Type-I interferon response during viral infection [18]. PTEN/Akt/PI3K signaling has previously been shown to maintain prostate cancer stem-like cell populations and the same study provided the gene expression data for the PC3 and DU145 sphere-forming and monolayer cultures used in Figure 1A [13]. Having identified the 14 genes in part using data from this study, we therefore went on to assess whether the expression of these genes is also PTEN-dependent. To do so, we knocked down PTEN expression in the DU145 cell-line and, having confirmed that this led to increased Akt phosphorylation (**Figure 2A**), we then used RT- PCR to assess the levels of the fourteen transcripts. We found significant increases in the expression of thirteen of the fourteen genes upon PTEN knockdown, the exception being IRF9 (**Figure 2B**). To assess the impact of restoring PTEN expression on the spherogenic potential of cells, we used a derivative of the PC3 cell-line in which wild-type PTEN was restored and induced upon Tetracycline treatment (TET). This led to the ablation of Akt phosphorylation (**Figure 2C**) and a significant reduction in spherogenesis by half (**Figure 2D and E**).

**Figure 1.**
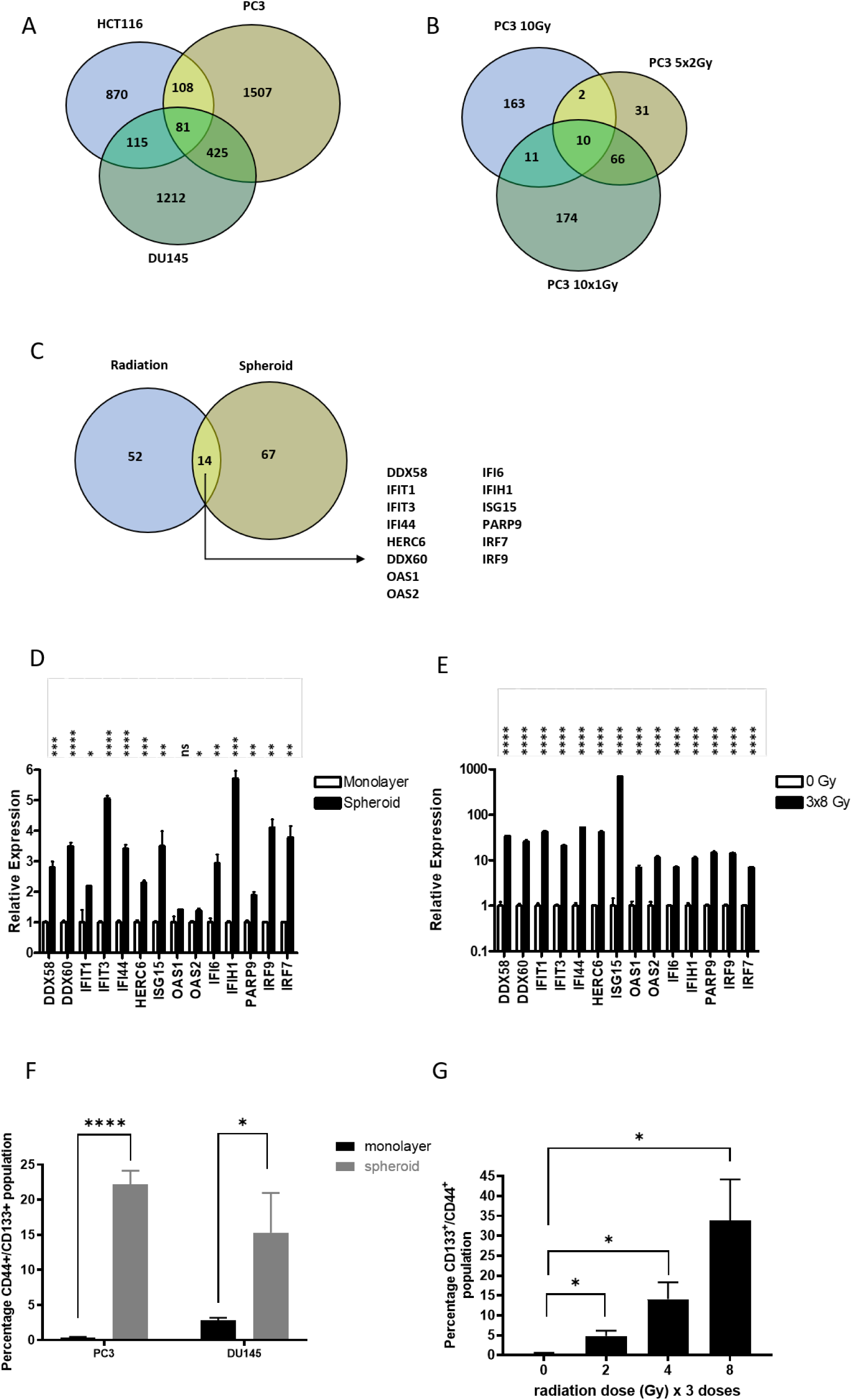
Meta-analysis of publicly available data identified conserved transcriptional characteristics of spherogenic and radiation-surviving cells. **A)** Publicly available microarray datasets for three cell lines (HCT116, DU145 and PC3) were analyzed and identified common upregulation of 81 genes following cell crowding/spheroid culture (Sphere). **B)** Publicly available microarray datasets of PC3 cells exposed to fractionated doses of radiation (5×2 gy and 10×1 Gy) or single high dose of radiation (10 Gy) were analyzed and identified common upregulation of 66 genes following multiple radiation doses. **C)** Common genes were identified to be upregulated by both cells crowding culture (Spheroid) and multiple radiation doses (Radiation). 14 commonly upregulated genes were identified: DDX58, DDX60, IFIT1, IFIT3, IFI44, HERC6, ISG15, OAS1, OAS2, IFI6, IFIH1, PARP9, IRF9 and IRF7. **D)** PC3 cells grown in monolayer and spheroid conditions were assessed for expression of the newly identified stress responsive transcripts. E) Expression of the newly identified stress responsive transcripts following 3 doses of 8 Gy radiation (72 hours post final dose). F) Induction of a ‘stem-like’ cell population (CD133+ and CD44+) in monolayer and spheroid conditions in both PC3 and DU145 cell lines, identified by flow cytometry. **G)** Radiation induced stem-like cell populations (CD133 and CD44 positive) in the PC3 cell line following multiple doses of radiation (3 times 2, 4 or 8 Gy). **D-G)** All experiments represent the average of at least 3 independent experiments, p values are * <0.05 **<0.01 ***<0.001 ****<0.0001.

**Figure 2.**
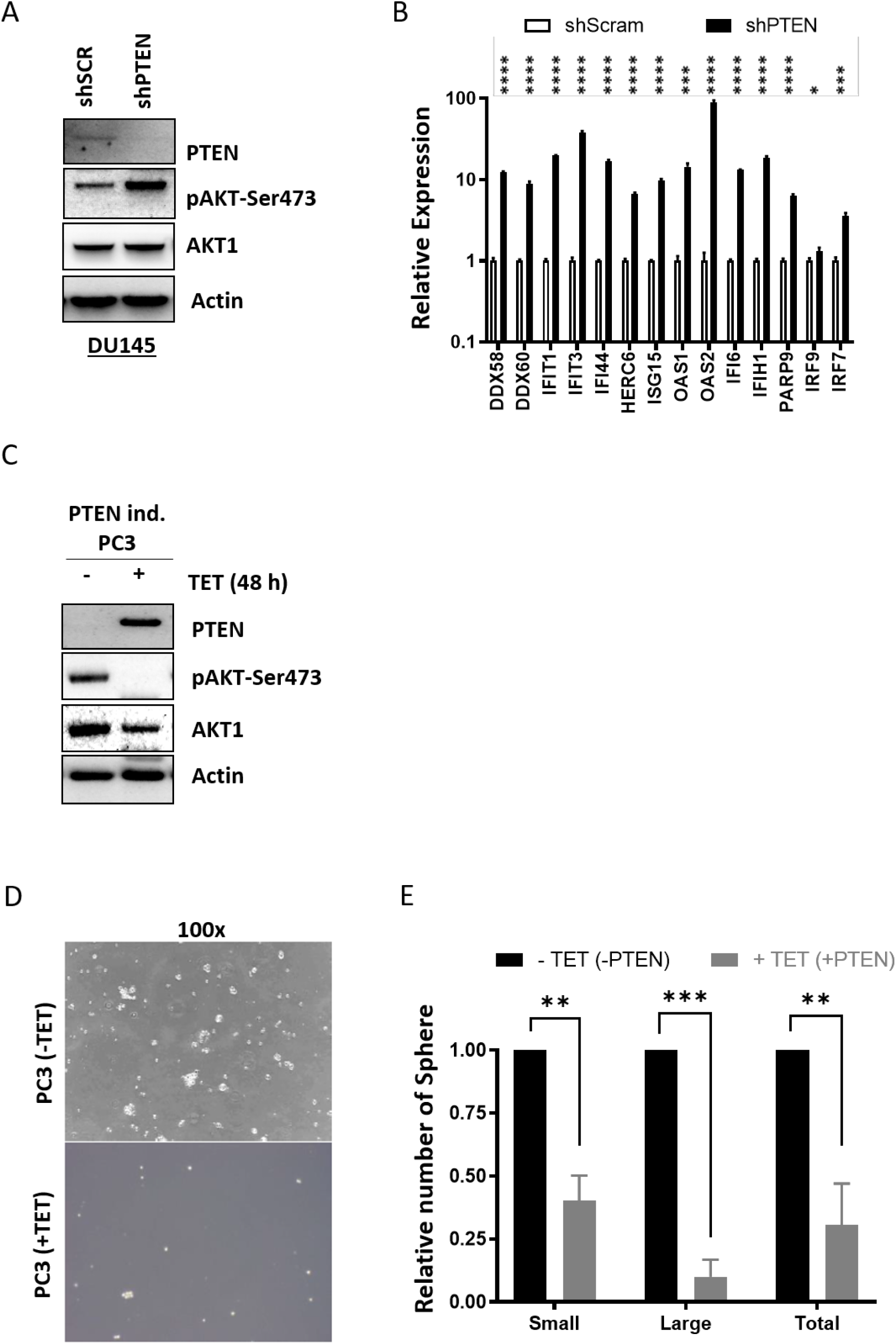
The expression of the newly identified stress responsive transcripts is PTEN dependent. **A)** PTEN knockdown in DU145 prostate cancer cells. Confirmation of PTEN knockdown and also activation of AKT phosphorylation, **B)** Assessment of stress responsive transcripts by real-time PCR in PTEN knockdown DU145. n = 3 independent experiments. **C)** Western blot validation of tetracycline (TET) inducible expression of PTEN in the PC3 cell line. Expression of PTEN suppresses AKT phosphorylation. **D)** PTEN expression suppresses the sphere forming potential of the PC3 cell line as quantified in **(E)**. All experiments **(A-E)** are expression of n = 3 independent experiments, p values are * <0.05 **<0.01 ***<0.001 ****<0.0001.

To elucidate the role of IRF7 as a mediator of cell survival in response to radiation, we further assessed the impact of different radiation regimens (i.e., 3×2 Gy, 3×4 Gy and 3×8 Gy) and spherogenic growth conditions on IRF7 expression and activity in PC3 cells. We observed an increase in both the mRNA and protein expression of IRF7 in cells surviving different fractionated exposures (**Supplemental Figure 2A**). IRF7 is known to translocate to the nucleus when activated by phosphorylation. Immuno-staining for IRF7 in cells treated with fractionated exposures revealed increased nuclear IRF7 (**Supplemental Figure 2B**). Increased nuclear IRF7 was also observed in cells cultured in sphere-forming conditions relative to monolayer cultures (**Supplemental Figure 2C**). To further confirm that increased nuclear IRF7 reflected changes in IRF7 phosphorylation we used a phospho-IRF7 specific antibody (phospho-Serine-477) for immuno-staining and found that this was increased in the nuclei of cells grown in sphere-forming conditions (**Supplemental Figure 2D**).

To further examine the role of IRF7 as the transcription driver, we overexpressed IRF7 in the PC3 cell line (**Supplemental Figure 3A**). IRF7 is known to shuttle from cytoplasm to the nucleus in response to phosphorylation by multiple kinases including IKKO and TBK1 [22] [23]. Overexpressing a phospho-mimetic IRF7 mutant [24], we enhanced the expression of 13 of the 14 transcripts we have newly identified (**Supplemental Figure 3B**). Also, the localization of the phospho-mimetic IRF7 mutant was primarily nuclear, unlike the wild-type IRF7, which is predominantly cytoplasmic in the unstressed state (**Supplemental Figure 3C-D**; 0 Gy vs 3×8 Gy).

To evaluate the clinical significance of IRF7 protein expression we stained samples from the Helsinki Prostate Cancer TURP tissue microarray [25]. By assessing the percentage of IRF7-positive nuclei (0-100) in luminal epithelial cells, we found that high nuclear IRF7 expression when combined with PTEN-loss, as determined by immunohistochemistry, was associated with reduced overall survival in patients (p = 0.029) (**Supplemental Figure 4A-D**, **Table 1**). We scored regions as areas of PTEN- loss based on PTEN staining that was negative or substantially reduced in cancerous prostatic epithelia as compared to adjacent benign prostatic epithelium (for more details, Material and Methods). Using an immunofluorescence approach to stain an independent cohort, the Salford prostate cancer TURP and needle-core biopsy TMA cohort (**Table 2**), we also observed an association between a high ratio of nuclear:cytoplasmic IRF7 staining in cancers and poorer prognosis for PCa patients. As described in material and Methods and Table 3, we calculated univariate HR: 1.82 (1.30-2.55), with p < 0.001; and multivariate HR: 1.51 (1.00-2.29), with p = 0.052) (**Supplemental Figure 4E and F**, **Table 3**). Immunofluorescent staining allowed us to accurately quantify the relative expression of IRF7 and PTEN in cancer regions versus in stroma and benign epithelial cells (**Figure 3A**). This approach revealed that high nuclear IRF7 expression alone in cancer regions is strongly associated with poor prognosis disease (univariate HR: 1.66 (0.98-2.80), p=0.057; multivariate HR: 1.37 (0.69-2.69), p=0.368) (**Figure 3B**, **Table 3**), low PTEN expression is also associated with poor prognosis disease as has previously been reported in the literature (univariate HR: 1.50 (1.07-2.09), p=0.018; multivariate HR: 1.38 (0.90-2.13), p=0.142) (**Figure 3C**, **Table 3**). Since we were able to multiplex these stains, we were also able to assess the power of combining the scoring for these markers in the same cell populations within the samples. We found that a combined score comprising high nuclear IRF7 expression and low PTEN expression defined the worst prognosis cases within this cohort (univariate HR: 2.46 (1.57-3.84) p<0.001; multivariate HR: 1.87 (1.04-3.37), p=0.037) relative to other combination scores for these two markers (PTEN+IRF7-, ref.; PTEN+IRF7+, univariate HR: 1.70 (1.11-2.60) p= 0.014/multivariate HR: 1.57 (0.95-2.61) p=0.079; PTEN-IRF7-, univariate HR: 1.33 (0.77-2.28) p= 0.301/multivariate HR: 1.49 (0.77-2.89) p=0.233) (**Figure 3D**, **Table 3**). The only combination of markers that was therefore significant when subjected to multivariate analysis was high nuclear IRF7 expression in samples with low PTEN expression.

**Figure 3.**
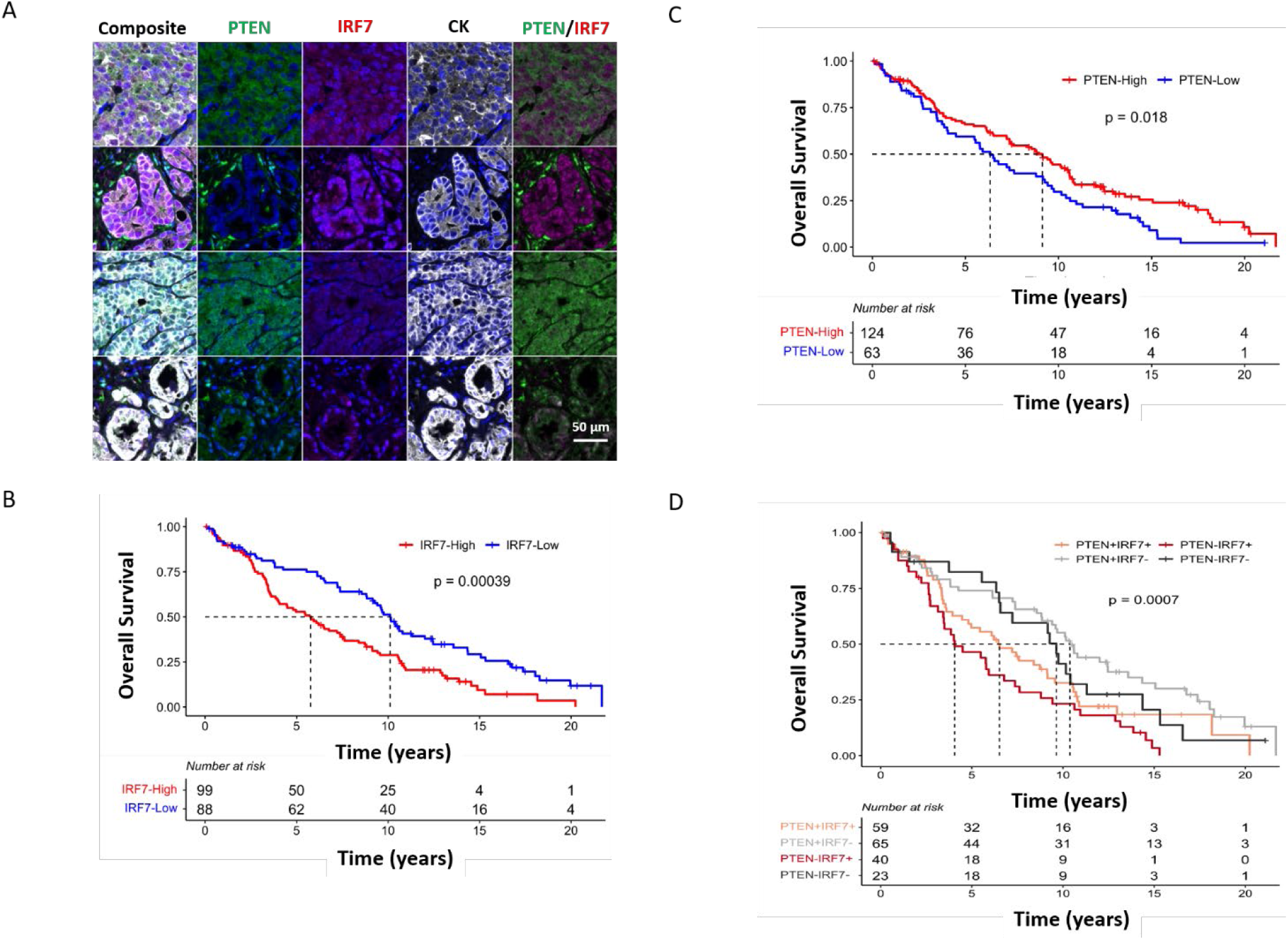
Increased nuclear IRF7 expression is prognostic in cases where expression is low or lost as determined by immuno-staining in the Salford prostate cancer TURP and needle-core biopsy TMA cohort. A) Representative images from Salford prostate cancer TURP and needle-core biopsy TMA. Patients with (B) nuclear IRF7 overexpression and (C) PTEN-loss in tumour cores had poorer overall survival. D) Patients with concomitant IRF7 overexpression and PTEN-loss (*IRF7^+^PTEN^-^,* in red, p= 0.0007) had poorer overall survival compared to the remaining cohort ((*PTEN+IRF7-,* ref.; *PTEN+IRF7+*, p= 0.014; *PTEN-IRF7-*, p= 0.301)). Dotted line denotes median survival. The most statistically relevant Log-rank statistic *p* values are reported in the figure.

**Table 1:**
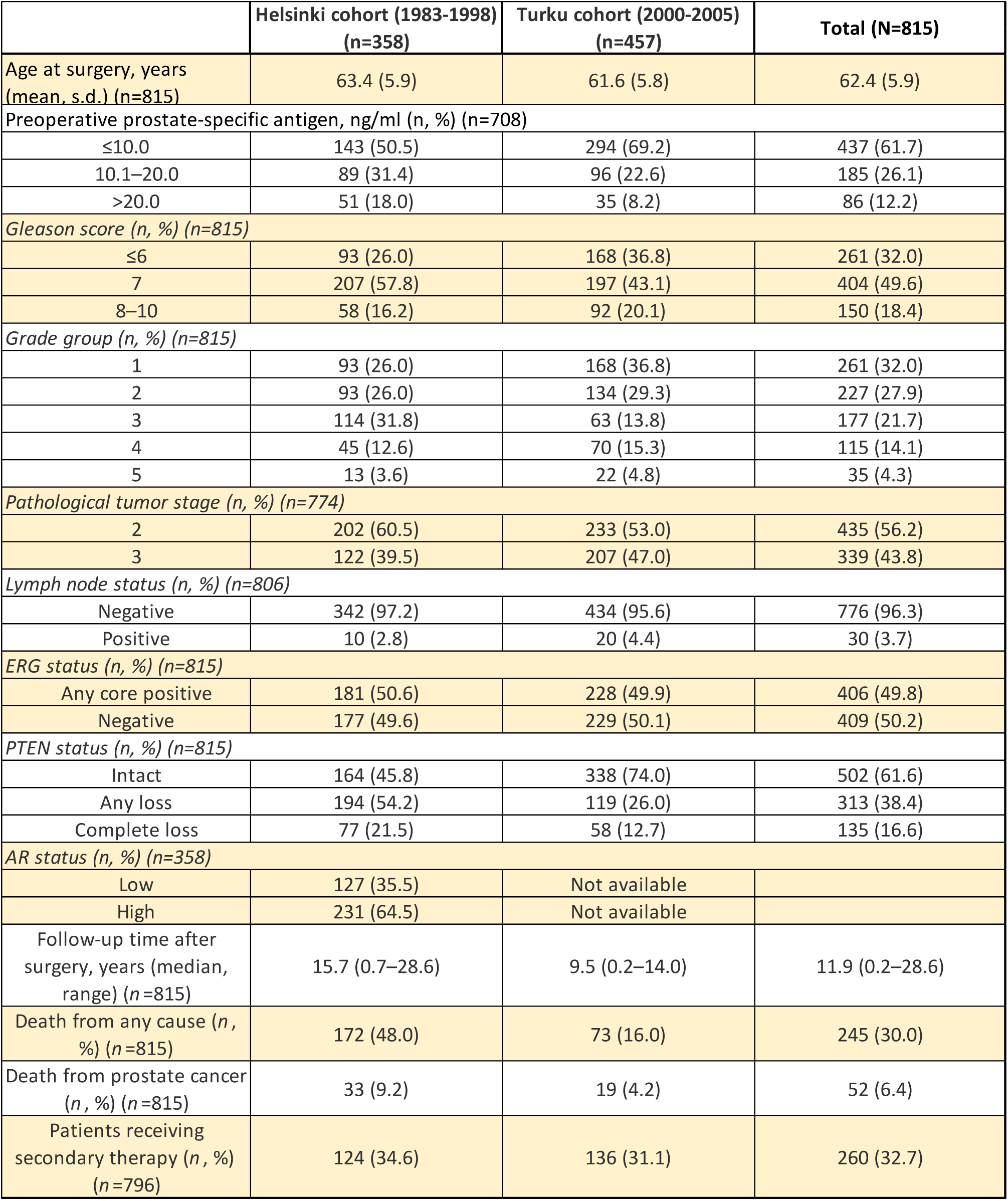
Clinical characteristics of patients included in the Helsinki Prostate Cancer TURP tissue microarray [25]

**Table 2:** Clinical characteristics of patients included in Salford TMA cohort. . Percentages reported in brackets, unless otherwise stated. IQR: Inter-quartile range

|  |  | **Total** |
| --- | --- | --- |
| **Total patients** |  | 187 |
| **Age at diagnosis** | Median (IQR) | 72 (11) |
| **Age group** | <65 years | 32 (17.1%) |
|  | 65-74 years | 73 (39.0%) |
|  | >74 years | 76 (49.6%) |
|  | Missing | 6 (3.2%) |
| **PSA at diagnosis** | Median (IQR) | 23 (49) |
| **PSA group** | 0-9.9 | 42 (22.5%) |
|  | 10-19.9 | 38 (20.3%) |
|  | 20-99.9 | 79 (42.2%) |
|  | >=100 | 24 (12.8%) |
|  | Missing | 4 (2.1%) |
| **ISUP grade group** | Groups 1-2 | 100 (53.6%) |
|  | Group 3 | 19 (10.2%) |
|  | Groups 4-5 | 29 (10.7%) |
|  | Missing | 48 (25.7%) |
| **Clinical tumour stage** | T1-T2 | 81 (43.3) |
|  | T3 | 67 (35.8) |
|  | T4 | 28 (15.0) |
|  | Missing | 11 (5.9) |

**Table 3:** Univariate and multivariate Cox regression models for Salford TMA cohort. Adjusted for age and PSA at diagnosis, Gleason grade group and clinical tumour stage.

| **OS** | **Category** | **N** | **HR (univariable)** | **HR (multivariable)** |
| --- | --- | --- | --- | --- |
| **IRF7** | low | 88 (47.1) | Ref | Ref |
|  | high | 99 (52.9) | 1.82 (1.30-2.55, p<0.001) | 1.51 (1.00-2.29, p=0.052) |
| **PTEN** | high | 124 (66.3) | Ref | Ref |
|  | low | 63 (33.7) | 1.50 (1.07-2.09, p=0.018) | 1.38 (0.90-2.13, p=0.142) |
| **PTEN/IRF7** | PTEN+IRF7- | 65 (34.8) | Ref | Ref |
|  | PTEN+IRF7+ | 59 (31.6) | 1.70 (1.11-2.60, p=0.014) | 1.57 (0.95-2.61, p=0.079) |
|  | PTEN-IRF7- | 23 (12.3) | 1.33 (0.77-2.28, p=0.301) | 1.49 (0.77-2.89, p=0.233) |
|  | PTEN-IRF7+ | 40 (21.4) | 2.46 (1.57-3.84, p<0.001) | 1.87 (1.04-3.37, p=0.037) |
| **IRF7** | low | 19 (10.2) | Ref | Ref |
| **Nuclear:Cytoplas** | high | 168 (89.8) | 1.66 (0.98-2.80, p=0.057) | 1.37 (0.69-2.69, p=0.368) |

Next, we demonstrated that the newly identified signature of 14 genes was all interferon regulated, including IRF7 (**Supplemental Figure 5A**). To gain more insight into the biological contribution of IRF7 to cancer, we generated stable IRF7 knockdowns in PCa cells (**Supplemental Figure 5**, **Figure 4**). IRF7 levels were significantly reduced compared to control cells by 90% and remained suppressed after interferon treatment (**Supplemental Figure 5B-C**). The interferon-stimulated gene factor 3 (ISGF3) complex (STAT1, STAT2 and IRF9) is a component of the IFN-triggered JAK/STAT pathway which transcribes genes, including IRF7 [26] [27]. Initially, this induces an acute innate immune response, that subsides into an adaptive DNA damage resistance-innate immune signal characterized by elevated STAT1/2 transcription to promote survival [28]. This complex is an established master regulator of the Type 1 interferon response. Its activity is regulated by phosphorylation and the majority of the newly identified 14 transcripts are known to be regulated by the unphosphorylated form of the complex, U-ISGF3 (**Supplemental Figure 5D**). Having established that the 14 genes are all IRF7- dependent (i.e., ‘IRF7-Sig’), we went on to blot for ISGF3 complex proteins in our knockdown and wild-type cells with and without radiation exposure. We observed a reduction of STAT1 and STAT2 expression in IRF7 knockdowns in response to irradiation (3×8 Gy, **Supplemental Figure 5E**), suggesting that IRF7 plays an important role in regulating the stability of this important transcriptional complex. Phenotypically, IRF7 knockdown transiently reduced proliferation (**Supplemental Figure 5F**). Nonetheless, after the initial delay in growth, IRF7 knockdown (shIRF7) cells recovered their proliferation, compared to scrambled controls (shscram) within 30 to 40 days (**Supplemental Figure 5F**), to then go into a growth crisis suggesting that a genetic knockout of IRF7 would not yield viable cells. The IRF7 knockdown cells had a cobblestone appearance compared to the fibroblastic phenotype of control cells (**Supplemental Figure 5G**). IRF7 knockdown significantly reduced spherogenic potential by almost 70% (**Figure 4A**) and associated numbers of stem-like cell populations by almost half, when the cells were grown under crowding conditions (**Figure 4B**). We achieved the a similar reduction (60%) in sphere formation when we restored PTEN expression in PC cells (Figure 2D-E) suggesting that IRF7 and PTEN regulate some shared biological processes. When knockdown and wild-type cells were treated with 3×8 Gy irradiation we observed a significant reduction in the number of surviving cells (**Figure 4C**). This inversely correlated with the levels of apoptosis as assessed by annexin V staining, which was significantly higher in the irradiated knockdown population compared to wild type (**Figure 4D**). The co-expression of markers of stem/progenitor cells, CD44 and CD133, was also significantly reduced in the surviving knockdown cells compared to wild type (**Figure 4E**). Collectively, these data suggest that IRF7 is necessary for the adaptive responses to stress, rather than as a prototypical oncogene driving cellular transformation. IRF7 is not itself a direct therapeutic target. However, as demonstrated, its activity is regulated by phosphorylation as is the activity of ISGF3, which creates opportunities to target the regulatory kinase activities instead. To test whether the inhibition of the stress-responsive pathways identified in this study could enhance radiation response, we tested the effects of IKKO/TBK1 inhibition, as these proteins are known to be activated by IFNβ and directly phosphorylate STAT1, a component of the ISGF3 complex [29]. Inhibitors of IKKO/TBK1 (BX795, **Supplemental Figure 6A**) and IkBα phosphorylation (Bay11-7085, **Supplemental Figure 6B**) were both effectively inhibiting the cell growth of both non-irradiated (yellow) and irradiated (5×2 Gy, green) cells that remained less affected. Inhibitors of the IKK/TBK1 pathways have been previously applied as alternative therapeutic approaches to tumours resistant to MEK inhibitors [29], suggesting that these pathways can compensate for each other. When we inhibited MEK with PD- 184352, we reduced the proliferation of both unirradiated and radiation-survivor cells by 50% with higher doses of 16 and 160 µM respectively (**Supplemental Figure 6C**). By contrast, PD-184352 combined with BX795 reduced the proliferation of the radiation survivors alone by up to 50% (**Supplemental Figure 6D**). Furthermore, the combination treatment additionally suppressed the expression of 13 out of the 14 stress-responsive transcripts of the ‘IRF7-Sig’ (**Supplemental Figure 6E**).

**Figure 4.**
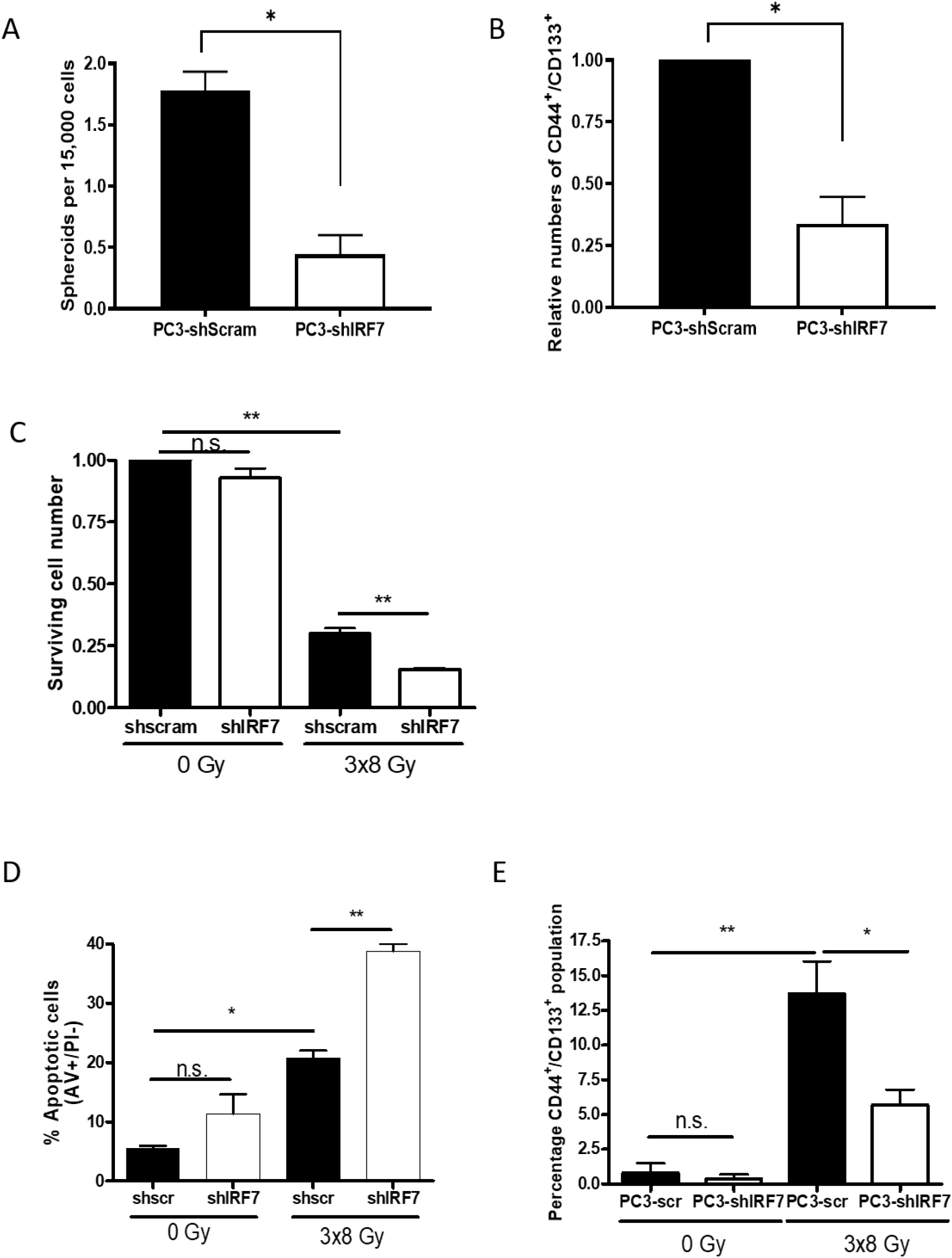
IRF7 modulates radiation response *in vitro*. **A)** Quantitation of the sphere forming capacity of PC3 cells shSram (expressing WT-IRF7 – black bars) and knockdowns for IRF7 (shIRF7 – white bars) assessed after 15 days of colturing. **B)** Normalised numbers of ‘stem-like’ cell populations (CD44^+^/CD133^+^) in PC3 control (shScram) and IRF7 knockdown (shIRF7) cells. **C)** IRF7 knockdown cells were more sensitive to multiple doses of radiation, as assessed by cell counts at 72 hours post radiation. **D)** Quantitation of apoptotic cells assessed by Annexin V/PI staining of shSram (expressing WT-IRF7 – black bars) and knockdowns for IRF7 (shIRF7 – white bars) after 72 hours in culture or after 72 hours form the last irradiation. **E)** Quantitation of the stem-like population ((CD44^+^/CD133^+^)) following multiple doses of irradiation (3×8 Gy) in PC3 cells shSram (expressing WT-IRF7 – black bars) and knockdowns for IRF7 (shIRF7 – white bars). All experiments **(A-E)** are expression of n = 3 independent experiments, p values are * <0.05 **<0.01.

The impact of IRF7 knockdown in response to irradiation was next assessed *in vivo*. Irradiated scrambled-control tumours (shScram) expressed IRF7 and displayed enhanced nuclear localization and phosphorylation at the endpoint (600 mm^3^ volume), compared to unirradiated controls (**Supplemental Figure 7A-B**), as was previously observed *in vitro*. Tumours grew at comparable rates with no significant difference in tumour volumes at the time of irradiation (**Supplemental Figure 7C**). However, following the treatment with 4 Gy, there was a significant delay in tumour re-growth (p = 0.0002) in the IRF7 knockdown xenografts, compared to the irradiated controls with re-growth commencing at ∼12 days post- irradiation in the wild-type and around 24 days post-irradiation in the knockdown (**Figure 5A-B**). To define the stable changes in the tumours following treatment we harvested tumours from knockdown and scrambled-control engraftments once they attained terminal volume. No significant differences in proliferation were observed between control and IRF7 knockdown tumours, as assessed by Ki67 staining (**Figure 5C**). Evaluation of the tumour microenvironment indicated that knockdown xenografts displayed reduced vascularization (mean vessel density – MVD - and CD31 expression) compared to the controls prior to irradiation (**Figure 5D**) and also reduced infiltration of CD11b-positive cells (**Supplemental Figure 7D**). This might be an effect mediated by interferon signalling that is known to impact the immune microenvironment during infection [18] [30]. These data imply that IRF7 knockdown primes an *in vivo* tumour microenvironment which supports a more durable response to radiation by influencing immune cell infiltration and tissue vascularity. To establish the impact of irradiation *in vivo* on the expression of the ‘IRF7-Sig’ transcripts, we went on to measure the levels of the 14 genes comprising the signature within wild-type tumour samples pre- and post-irradiation using RT-PCR (**Figure 5E**). All these transcripts, with the exception of IFIT1, were increased post-irradiation (**Figure 5E**). Next, we assessed whether the levels of these transcripts were affected in the IRF7 knockdown tumours versus wild-type post-irradiation (**Figure 5F**). IRF7 knockdown was associated with reduced expression of all the ‘IRF7-Sig’ genes post-irradiation relative to wild-type tumours with the greatest impact being on OAS1 and OAS2 (**Figure 5F**). These *in vivo* data support further evidence the IRF7 dependency of these genes.

**Figure 5.**
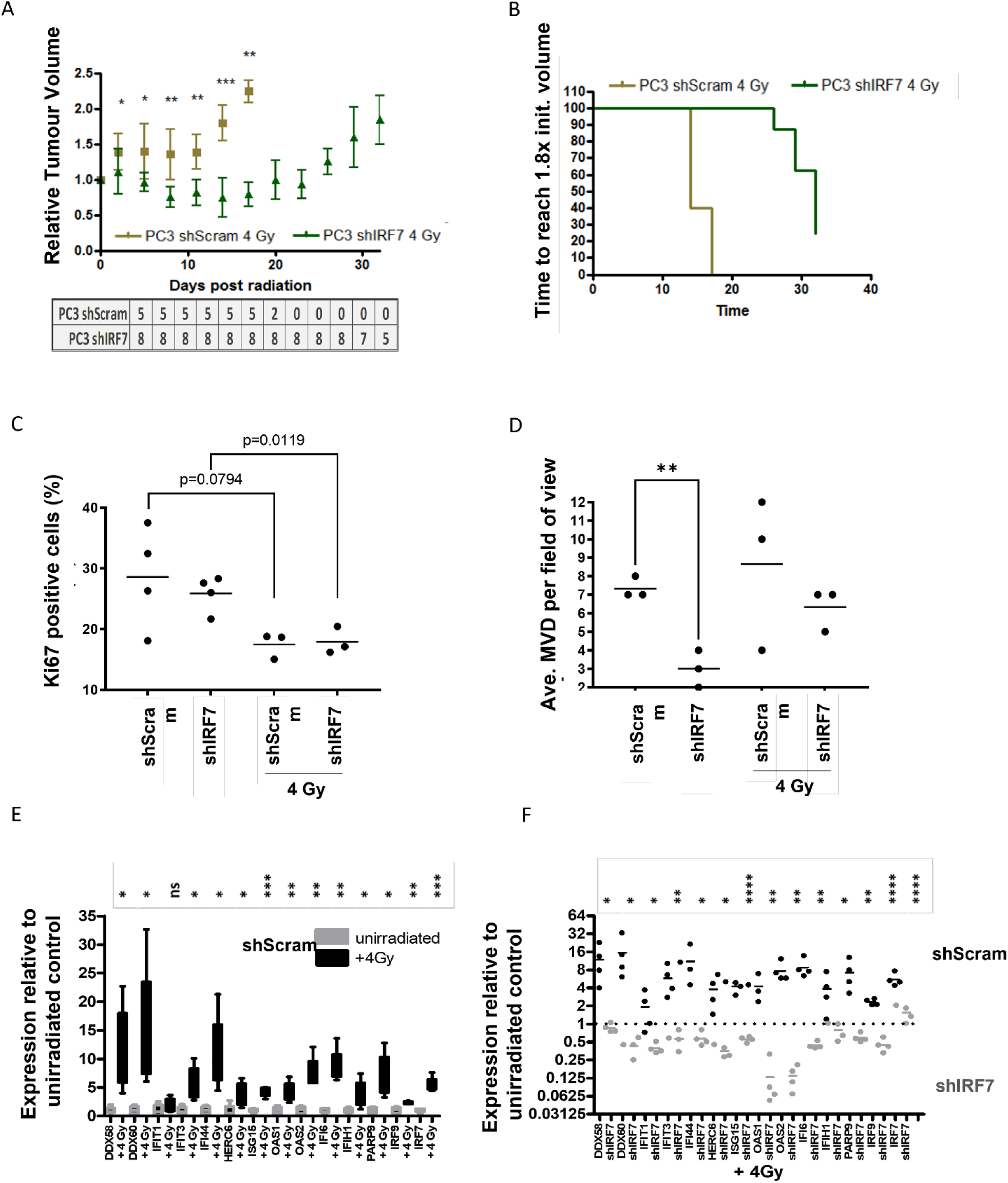
IRF7 modulates radiation response *in vivo.* **A)** Response of IRF7 knockdown xenografts to radiation. Tumours were grown to approximately 200-300 mm^3^ before receiving a single dose of radiation. Numbers of animals included in the analysis are shown below the graph. p values are * <0.05 **<0.01 ***<0.001. **B)** Graphical representation of the time in which the tumours reached maximal volume. **C)** Proliferation (Ki67) and **(D)** vascularisation (CD31^+^) were assessed in unirradiated xenografts of control and IRF7 knockdown PC3 cells and quantified. **E)** Assessment through RT-PCR of stress responsive transcripts in harvested tumours at the end point in shScram- derived tumours unirradiated (grey bars) and following 4 Gy irradiation (black bars). Expression is normalised to unirradiated tumours. **F)** Assessment through RT-PCR of stress responsive transcripts in harvested tumours at the end point both in shScram- derived tumours (black dots) and in shIRF7- derived tumours (grey dots) following irradiation (4 Gy). **D-F)** p values are * <0.05 **<0.01 ***<0.001 ****<0.0001.

To more comprehensively define the transcriptional response to irradiation and the contribution of IRF7 to this response, we generated RNA-Seq data from wild-type PC3 cells and IRF7 knockdown PC3 cells that were either mock irradiated or radiated with 1, 2 or 3 doses of 8 Gy. We identified 6290 significantly altered genes across the four treatment groups (p<0.05), with ‘IRF7-Sig’ upregulated in the irradiated controls (IRF7 wild type, top right quadrant **Figure 6A**) relative to IRF7 knockdown (left quadrants **Figure 6A**). This evidence further confirmed IRF7 as a regulator of the expression of ‘IRF-Sig’. To gain further insight into the role of IRF7 in response to irradiation of PCa cells, genes with a fold-change magnitude greater than 1.5 were retained and sorted into four separate gene lists: IRF7 knockdown irradiated (n=535), control irradiated (n=819), IRF7 knockdown untreated (n=25), and control untreated (n=80). All genes were filtered against the HumanNet network [31], and the four condition-specific gene lists were merged to create the ‘combined’ gene list. These five gene lists were analysed with NetNC [32]. Our ‘comprehensive’ network highlights biological processes including viral immune response, proliferation (nucleosome assembly) and keratins (Supplemental Figure 8). Immunological processes were only identified by NetNC under conditions of irradiation in the IRF7-expressing (control, irradiated) condition. Results also showed that the irradiated control (IRF7 wild type) network contained 13 genes of the 14 genes- ‘IRF7-Sig’ (**Figure 6B**; **Supplemental Figure 9**). By contrast, irradiated IRF7 knockdown cells had clusters containing keratins and other related pathways associated with cell adhesion and cell differentiation (**Figure 6C**; **Supplemental Figure 10**). Network analysis indicated druggable targets (**Table 4** – drug interaction groups 3 -, **Supplemental Figure 11**). Single strong candidate targets were identified for both the wild-type irradiated (IDO1, ellipse-shaped node in **Figure 6B** and **Supplemental Figures 8 and 9**) and the knockdown irradiated networks (XPNPEP2, ellipse-shaped node in **Figure 6C** and **Supplemental Figure 10**). Given the increased sensitivity to radiation that we have already shown in IRF7 knockdown cells we went on to validate IDO1, the predicted druggable candidate target for wild-type irradiated conditions, and a metabolic enzyme that can modulate the tumour immune micro-environment [33]. We confirmed that IDO1 expression is significantly increased in prostate cancer cells surviving radiation (3×2 Gy and 3×8 Gy, **Supplemental Figure 12**). Epacadostat is a selective IDO1 inhibitor [34] identified in our network analysis. We treated PC3 cells that had survived multiple doses of radiation with Epacadostat (0.1, 1, 10 and 50 nM), leading to effective cell killing of 30% and 90% of the cells surviving 3×2 Gy (green) and 3×8 Gy (red) respectively within 72 hours from the last dose (Figure 6D, p < 0.001). The effect also correlated with the levels of IDO1 mRNA induced by the radiation with cells surviving 3 doses of 8 Gy being the most sensitive to the inhibition (**Figure 6D**, **Supplemental Figure 12**). These data together suggest that IRF7-dependent target genes, including IDO1, are part of a stress-responsive biology that supports cell survival upon irradiation. Indeed, we have also shown that targeting either kinases upstream of IRF7, or the IRF7 target gene IDO1, can effectively diminish the number of cells surviving radiotherapy.

**Figure 6.**
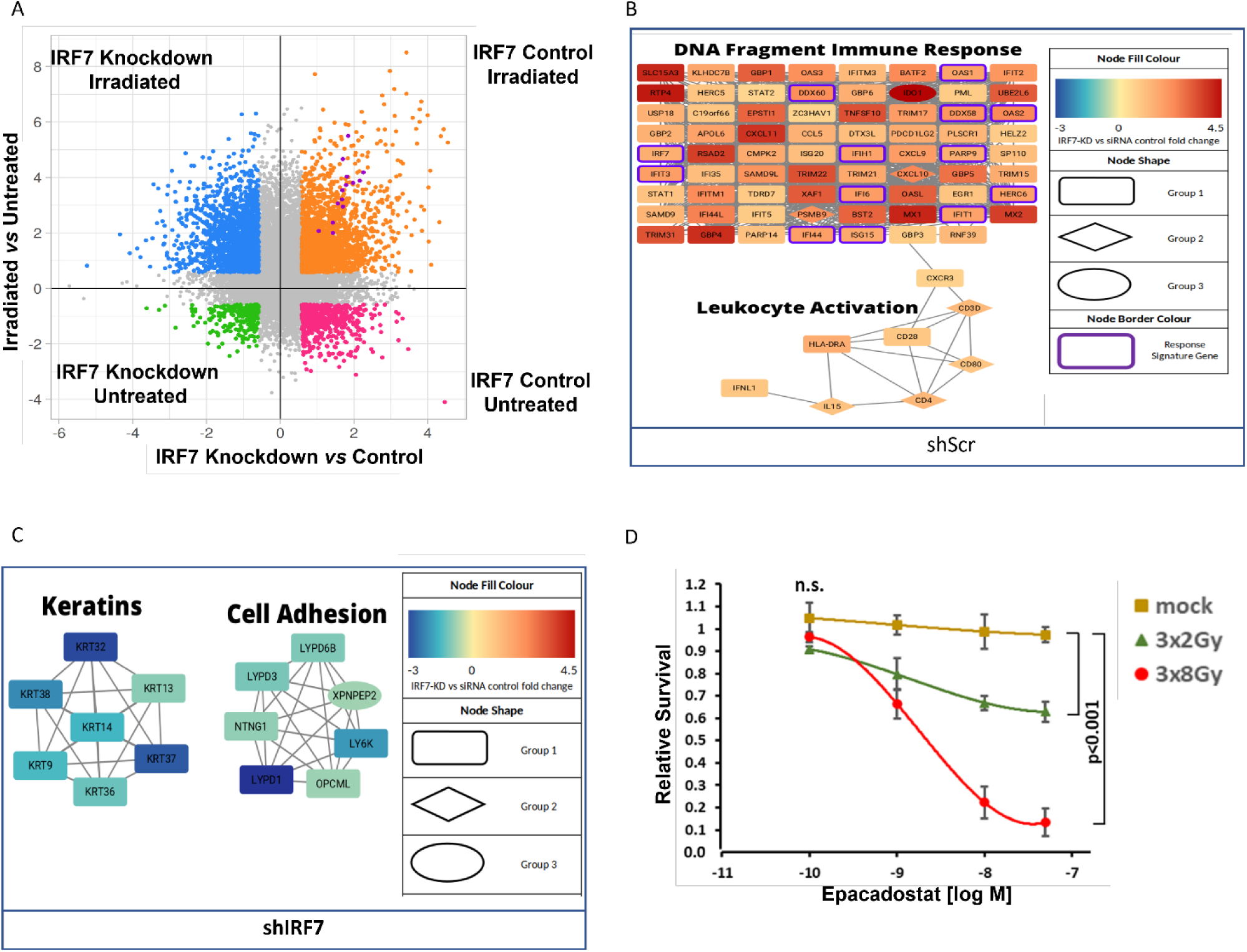
Network analysis identifies candidate druggable targets associated with IRF7-dependent radiation response. **A)** Points represent log2 fold-change values calculated across two axes: irradiated versus untreated and IRF7 versus control, forming four quadrants. Thresholds for 1.5- fold change in each axis specify four sectors (blue, orange, green, pink) which defined the four gene- lists taken forward into NetNC analysis. The IRF7-Sig genes (purple) are all found within the irradiated, IRF7 wild-type sector (orange). **B) and C)** show key clusters from the irradiated IRF7 wild- type (panel B) and irradiated IRF7 knockdown) (panel C) networks; respectively representing output of NetNC analysis from orange and blue genes shown in panel A. Node colour shows fold-change with respect to IRF7 status (x-axis coordinates in panel A) from highest expression in IRF7 wild-type (dark red) to highest expression in IRF7 knockdown (dark blue). Thirteen of the IRF7-Sig genes (purple border) are present in the IRF7 wild-type network. Results from network druggability analysis are indicated by node shape for candidate targets (diamonds), high-scoring candidate targets (ovals) and other genes (rectangles). Network cluster annotations are representative of functional enrichment results. **D)** Validation of the high-scoring candidate target IDO1 and candidate drug Epacadostat from the result shown in panel B. PC3 cells treated with Epacadostat (0.1,1, 10 and 50nM) for 72 hours after multiple doses of irradiation; mock/not radiated (yellow), 3×2 Gy (green) and 3×8 Gy (red). The mean of three independent experiments is shown. Treatment with Epacadostat causes a significant reduction in PC3 cell survival at both radiation doses relative to the control (p<0.001).

**Table 4:** Prioritisation criteria and candidate drug target groupings applied to the Network Analysis.

| Drug Interaction Group | Description | WT_irr Main Cluster Genes | KD_irr Full Network Genes |
| --- | --- | --- | --- |
| **0** | All condition-specific network genes | 79 | 34 |
| **1** | No known/predicted drug-gene interaction | 64 | 23 |
| **2a** | No ChEMBL interaction record | 12 | 8 |
| **2b** | Does not satisfy Lipinski's "Rule of Five" | 1 | 1 |
| **3a** | Potential drug candidate<br>Limited impact on network | 1 | 1 |
| **3b** | Recommended drug candidate | 1 | 1 |
|  | Greater impact on network |  |  |

## Discussion

It is well established that selection for stem-like populations contributes to radiotherapy resistance, tumour recurrence and subsequent disease progression following radiotherapy treatment Wang [35] [36]. Our findings have identified a Type-I interferon driven stress-responsive pathway that is upregulated in response to multiple cellular stresses, including radiation in PCa. We have demonstrated that IRF7 is a key mediator of this stress-responsive signature and is required for the occurrence of stem-like populations in response to low-dose fractionated radiation *in vitro* and *in vivo*. Cancer stem cells have been often described as tumour initiating cells Rajasekhar [12] and have been extensively reported as contributors to the acquisition of radiotherapy resistance Arnold [14] and metastasis formation [37]. We believe subpopulations of surviving stem-like cells contribute to radiotherapy and multi-drug resistance. In this study we have shown that IRF7 can support both the expression of a network of stress responsive Type1 interferon genes and also the survival of cells treated with radiation. In addition, at the protein level the nuclear localisation of IRF7 in primary prostate cancer is a prognostic marker when assessed alongside PTEN expression. The latter observation strongly suggests that IRF7 activity and nuclear function is an important parameter to consider, and perhaps to target, to improve treatment response. From the literature we know that IRF7 nuclear translocation depends on its phosphorylation by a number of kinases including MEK, Akt and IKK. We have shown using a phospho-mimetic mutant of IRF7 than we can recapitulate the IRF7-dependent biology in prostate cancer cells. The knowledge that its activity depends on phosphorylation led us to assess the combinatorial impact of MEK and IKKε/TBK1 inhibition on cells surviving irradiation. We found enhanced sensitivity to a combination of BX795 and Bay11-7085 (inhibitors of IKK/TBK1 and IkBα respectively) in cells that had survived irradiation, relative to un- irradiated cells. This suggests that targeting these kinases or other radiation-induced IRF7- dependent druggable targets, may be therapeutically beneficial in preventing stress-resistant cell populations from emerging following radiation treatment, to ultimately improve outcome for PCa patients. We have also used other approaches to identify actionable targets in cells surviving irradiation. For example, our network analysis on the RNA-Seq data from IRF7 wild-type and knockdown cells identified IDO1 as a druggable target amongst Type 1 interferon response genes. IDO1 as a metabolic enzyme has important cancer cell-intrinsic and extrinsic effects arising from the production of kynurenine from tryptophan [38]. Cancer-cell intrinsic effects of IDO1 center on the DNA repair capacity of cancer cells. Kynurenine production provides an endogenous source of NAD+, which is a cofactor for PARP1 in DNA repair [39]. Consequently, levels may sustain the DNA repair capacity of cancer cells through maintenance of the available pool of redox cofactors. In addition, a recent study has shown that kynurenine is required to maintain the activity of a deacetylase, SIRT7, in glioma. SIRT7 activity has been reported to create a permissive chromatin landscape for 53BP1 recruitment to sites of DNA damage, supporting efficient DNA repair, by reducing Histone H3 lysine-18 acetylation (H3K18) at DNA damage sites [40]. The Epacadostat effects that we have observed may therefore reflect an impact on the DNA repair capacity of the treated cells, although the impact of kynurenine on these biologies remains to be explored in prostate cancer. For example, IDO1 has previously been described as responsible of the interferons triggered dormancy in tumour-repopulating cells and a driver of the immune suppressive state in PTEN deficient PCa tumours [41] [42]. Although IDO1 inhibitors have failed in clinical studies when combined with immunotherapy or radiotherapy, here we show that an IDO1 inhibitor is effective on cells that have survived radiation. Given that the same IRF7-Sig is induced in vivo we believe the same approach should be tested in in vivo models of tumorigenesis – pre-treatment with radiotherapy and subsequently treatment with an IDO1 inhibitor. The biological impact of IDO1 inhibition may be different *in vivo* versus *in vitro* because IDO1 activity affects both intrinsic tumour cell biology and also the tumour micro-environment. The effects of IDO1 inhibition would beneficial in both cases. IDO1 metabolizes tryptophan into kynurenine, which can suppress T-cell proliferation and promote regulatory T-cell development thus promoting the transition towards an immune suppressive tumour microenvironment [43]. An elevated tryptophan metabolism has also been described in relation to mechanisms of multi- drug resistance, through the activation of AhR/c-Myc/ABC-SLC transporters signaling pathway in PCa [44]. Therefore, the inhibition of IDO1 may restore T-cell function and reduce accumulation of regulatory T cells (extrinsic) but also reduce the activation of other therapy-resistance mechanism (intrinsic). Other genes identified within our network analysis, as upregulated in the IRF7 knockdowns following irradiation, pertained mostly to keratins and other related pathways associated with cell adhesion and cell differentiation. The genes within the IRF7 knockdown networks are functionally similar with many others known to be contributing to the extracellular adhesion of cells. This is the case for the keratins that have been shown to moderate apoptosis in specific conditions and protect epithelial cells from mechanical stress [29]. For example, a shortage of KRT14 has been shown to make cells more likely to undergo apoptosis [30]. Also, KRT6A, which has been shown to suppress cell viability and invasion [31], was instead upregulated in the IRF7 knockdown condition following irradiation. The network also includes LY6K, which has been shown to cause increased TGF-β signaling in breast cancer, contributing to cell growth, metastasis and EMT [33] Dhiman [34]. LYPD3 has been shown to contribute to tumorigenic development in metastatic melanoma [35]. Altogether these results suggest that although the knockdown of IRF7 achieved a greater apoptotic response to irradiation, the surviving cells may be sustained by the activation of resistance mechanisms susceptible to other targeting strategies.

Our study suggests that this IRF7-dependent adaptive pathway which would normally be activated in response to viral infection, is upregulated in PCa cells in response to treatment. We demonstrated that high nuclear levels of IRF7 associated to PTEN-loss are together indicators of poorer prognosis. This gives more insights on the possibility to utilize IRF7 as a predictive and prognosticating marker for prostate cancer patients also in relation to the staging of the disease and whether the specimen is derived from the primary tumour or the bone metastasis. In fact, IRF7 overexpression has been also reported to have protective implications for bone metastasis by increasing the IFN-beta-mediated NK cells activity [45]. Moreover, given the importance of evaluating whether the localization of IRF7 is nuclear or cytoplasmic, approaches involving bulk RNA-Seq for predictive markers might therefore not be as effective. In conclusion IRF7 expression and nuclear localization predicts poor survival in independent PCa patient cohorts, indicating that this biology is of clinical significance and requires further prospective clinical validation.

## Materials and Methods

### Cell culture

Cell lines were maintained in RPMI containing 10% FBS and penicillin and streptomycin, except for RWPE1 which were maintained in Keratinocyte Serum Free Medium supplemented with bovine pituitary extract and human recombinant epidermal growth factor (manufacturer supplied, Life Technologies). For spheroid cultures, cells were seeded at 2×10^6^ per 90mm plate coated with 20 mg/mL poly-HEMA, in medium defined in **Table 5** for 14 days. Spheres were counted manually and by cell sizing (Beckman Coulter, >40 µm). When cells were irradiated all cell lines were maintained in RMPI supplemented with 10% FBS. Medium was replaced immediately prior to each radiation dose. Clonogenic assays were performed as previously reported [46]. For each experiment, unexposed controls were prepared and treated as sham exposures (referred as mock or unir) and harvested at matched time intervals. For interferon alpha and beta (Peprotech) were used at the indicated doses and time. Adherent cells surviving radiation treatment were counted following trypsinisation (Beckman Coulter). Apoptosis was assessed by Annexin V labelling and propidium iodide exclusion (BD Biosciences). Inhibitors used in the study were supplied as follows: Docetaxel (Belfast Health and Social care trust), Bay 11-7085 (Santa Cruz Biotechnologies), BX795 (Invivogen), Dexamethasone and MEK inhibitor PD-184352 (Sigma-Aldrich). Cell viability assays were performed as previous reported Pickard [46]. dsRNA treatment was performed by transfecting 5’-ppp-dsRNA (Invivogen) at a final concentration of 0.5 µg/mL using RNAiMax (Life Technolongies) [47]. DU145 with PTEN knockdown were kindly provided by the laboratory of Professor David Waugh, QUB.

**Table 5:**
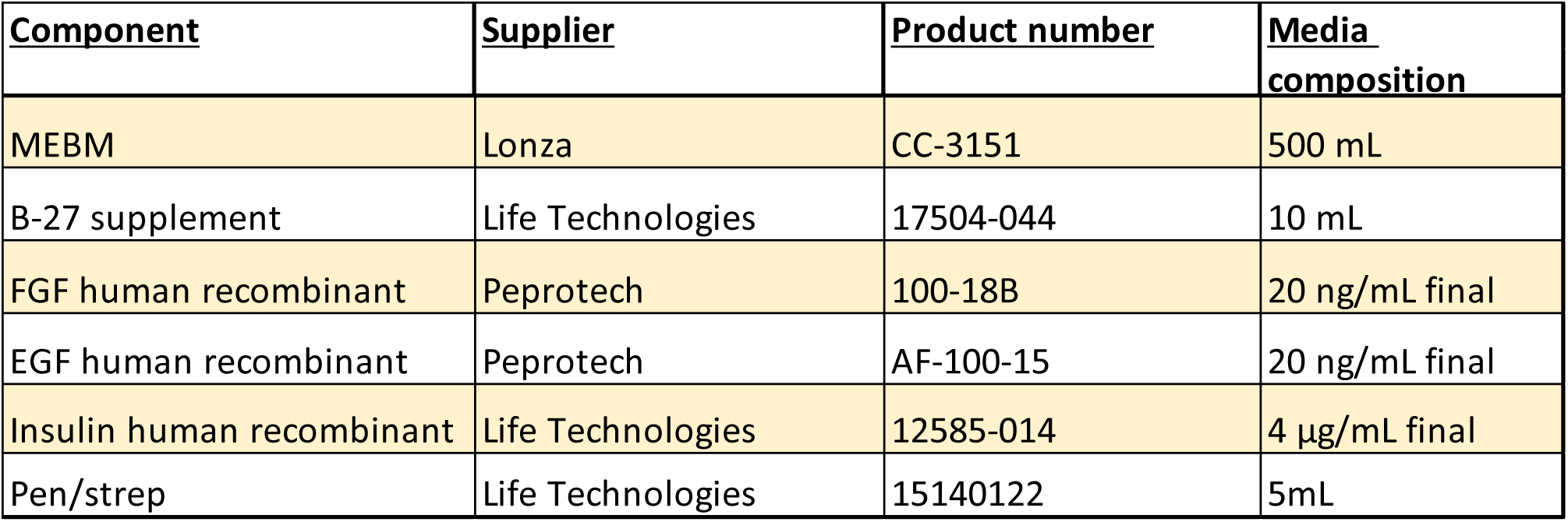
Components of spheroid culture medium.

### Constructs

shRNA to target IRF7, sequence: GGAGAAGAGCCTGGTCCTG, was taken from the retroviral shRNA NKI library [48]. Retrovirus generation was conducted as previously described Pickard [49]. Flag-tagged IRF7 and phospho-mimetic mutant were purchased from GenScript Biotech Corporation (Piscataway, NJ, USA).

### Flow cytometry

Flow cytometry was conducted as previously described in [13] using CD44-PE (BD biosciences) (1:100 dilution) and CD133-viobright FITC or CD133-APC (Miltenyi Biotec) (1:100 dilution).

### Immunofluorescence

For immunofluorescent detection of proteins, initially cell lines were fixed with 4% paraformaldehyde. Cells were permeabilised with 0.2% triton/PBS. After blocking (5% FBS/PBS) cells were stained overnight with the indicated antibodies. IRF7 (Santa Cruz) 1:100, Phospho-IRF7 (Ser 477) (Cell Signaling) 1:400. More specifications on the antibodies are listed in **Table 6.**

**Table 6:** List of antibodies utilised in this study.

| Protein | Antibody details |
| --- | --- |
| IRF7 | SCBT: sc-9083 (Rabbit), (WB: 1:200, IF: 1:200)<br>sc-74472 (WB: 1:200, IHC: 1:20) |
| GAPDH | SCBT: sc-25778 (WB: 1:2000) |
| STAT1 | SCBT: sc-417 (WB: 1:1000) |
| STAT2 | SCBT: sc-476 (WB: 1:1000) |
| IRF9 | Atlas antibodies: HPA001862 (WB: 1:500) |
| Phospho-IRF7 (Ser 477) | Cell Signaling: #12390 (WB: 1:500, IF: 1:400) |
| Beta-actin | Sigma: A2228 (WB: 1:10,000) |
| J2 (dsRNA) | Scicons: J2 (IF: 1:400) |
| Flag (M2) | Sigma: F1804 (IF: 1:400) |
| CD31 (PECAM1) | Aviva Systems Biology: #OAAB00949 (IF: 1:200) |
| CD11b | Abcam: EPR1344 (IHC: 1:4000) |
| Ki67 | Thermo Scientific RM-9106-S (IF: 1:400) |

### Immunohistochemistry and TMA

Study Population: A TMA of all RP treated patients treated in the University of Helsinki between 1983 and 1998 was utilized [25]. Freshly cut 4 μm thick TMA sections were mounted on slides and stained manually. Heat induced epitope retrieval was performed utilizing a buffer of Citric acid, pH 6.0. IRF7 antibody (SC-74472, Santa Cruz) was utilized at a concentration of 1:20. Slides were then digitized using a slide scanner with a 20X objective (Pannoramic 250 Flash III, 3DHistec, Budapest, Hungary). Slides were uploaded to a web-based browser (Webmicroscope, Fimmic Oy, Helsinki, Finland). IRF7 scoring was performed by assessing the percentage of positive nuclei (0-100) in luminal epithelial cells (AE and TM). Spot-wise scores from each spot per patient were pooled, to form mean IRF7 expression per patient. PTEN values were obtained from the same cohort, as described previously [50]. Briefly, PTEN was considered lost, if the PTEN expression intensity was negative or substantially reduced in cancerous prostatic epithelia as compared to adjacent benign prostatic epithelium. The expression of PTEN WT (e.g., full expression) is referred as Intact. Statistical analysis was performed using SPSS. Mean IRF7 staining in cancer cores, was 4.19 in Gleason ≤ 6, 5.78 in Gleason 7, and 5.65 in Gleason ≥ 8. In Univariate Cox Regression Analysis, patients with PCa which had any PTEN loss and IRF7 positivity had shorter Disease-Specific Survival after operative treatment. Full description of the Helsinki cohort is in **Table 1**.

The Salford prostate TMA [51] consisting of diagnostic needle core biopsies from 130 patients and TURP samples from 150 patients attending the Urology clinic of Salford Royal NHS Foundation Trust was covered by MCRC Biobank Ethics 10_NOCL_02, Manchester, UK. Patient characteristics are defined in Table 3. PCa tissue from these cores was identified and Gleason graded by a consultant histopathologist. Where available, normal-adjacent tissue cores were also identified. Median follow-up for the TMA cohort was 16.7 years. Full description of the Salford prostate TMA cohort is in **Table 2.**

Freshly cut 4μm TMA sections were mounted on slides, dewaxed and heat induced epitope retrieval was performed using a buffer of TRIS/EDTA, pH 9.0. Slides were incubated in IRF7 antibody (1:100, sc-74472, Santa Cruz) overnight at 4°C at a concentration of 2ug/ml. The slides were washed with PBS and wet loaded onto the Ventana Discovery Ultra automated IHC/ISH research platform. Following an HRP conjugation step a TSA Opal 570 1:150 (Tyramide Signal Amplification Opal 4-Color Manual IHC Kit, Perkin Elmer) was linked to the IRF7 antibody previously bound to the slide overnight. All further primary and secondary antibodies were bound to the slides and stained in a fully automated manner in this order, with a heat denaturation step at 95°C for 4 min in between: PTEN (D4.3) 1:50 (#9188 Cell Signalling Technology) / TSA Opal 520 1:100; pan cytokeratin (C11) 1:10,000 (C2931 Sigma-Aldrich/Merck) / TSA Opal 690 1:150; DAPI. Slides were mounted in ProLong Gold antifade reagent (Invitrogen) and scanned on a Perkin Elmer Vectra using the x20 magnification lens. Automated image analysis was performed using InForm v2.4 (Perkin Elmer). Operator screening was used to provide quality control for the automated core detection and region of interest (ROI) identification.

Data for 187 patients with non-metastatic PCa were available for survival analysis. Segmented single cell data were analysed on R Studio v1.1.423. Median epithelial IRF7 fluorescence in the nucleus and epithelial PTEN expression across the entire cell amongst all cells from tumour cores from individual patients were calculated. Proportion of cells with median epithelial IRF7 expression greater than the median expression within epithelial cells from normal-adjacent cores was used to determine IRF7 status. More specifically, the IRF7 nuclear:cytoplasmic ratio was calculated at a single cell level. Log10 transformed median N:C ratio was calculated across all tumour’s epithelial cells for each individual patient. Patients with median N:C> 0.6 were classified as ‘high’. Nuclear IRF & expression amongst tumour epithelial cells were compared against median epithelial cell expression amongst all cores from normal-adjacent regions. This classified individual cells as high/low IRF7 expression. Patients with > 55.5% IRF7 highly expressing cells were classified as IRF7-high. PTEN expression across tumour cores were normalised against paired normal-adjacent cores. Where paired normal-adjacent cores were unavailable, epithelial PTEN expression was normalized against adjacent stromal PTEN expression. PTEN status was determined as previously described Ahearn [52].

The survminer package was used to determine optimum thresholds to categorise patients with either “low” or “high” PTEN and IRF7 expression. Log-rank statistic p values are reported. Univariate and multivariate analysis of PTEN and IRF7 status and the correlation with survival are shown in **Table 3**.

### Datasets

Microarray datasets exploring the effects of spheroid culture conditions for PC3, DU145 and HCT116 were the following datasets: GSE10832 (PC3 and DU145) [13] and GSE53631 (HCT116) Zhang [17]. Differential gene expression in response to radiation was also examined in PC3 cells exposed to single or multiple radiation doses, GSE36720 (PC3 and DU145) [21].

### Xenograft

Experiments were performed using 4–5-weeks-old female Severe Combined Immunodeficiency (SCID) mice (Charles River Laboratories). PC3 cells expressing wild type IRF7 (shScram) and knockdown IRF7 (shIRF7) (2×10^6^) were implanted subcutaneously. Tumours were irradiated at a volume of approximately 200-300 mm^3^ with a dose of 4 Gy using a small animal radiotherapy research platform (SARRP, Xstrahl, UK) **(Supplemental Figure 7C)**. Treatments were delivered under cone-beam-CT image guidance with a parallel-opposed beam geometry at a dose rate of2.8 Gy/min. Tumours were measured every 3 days using callipers until they reached a target volume of 1.8x pre-irradiation volume or a maximum volume of 600 mm^3^. Tumour specimens were excised post-mortem and either kept at − 80 °C for RNA extraction or fixed in 1% Paraformaldehyde (Sigma Aldrich) for paraffin embedding.

### Antibodies

The antibodies utilised in this study are summarised in **Table 6**. **Real time PCR**

The primer sets utilised in this study are summarised in **Table 7**. **RNA sequencing**

**Table 7:**
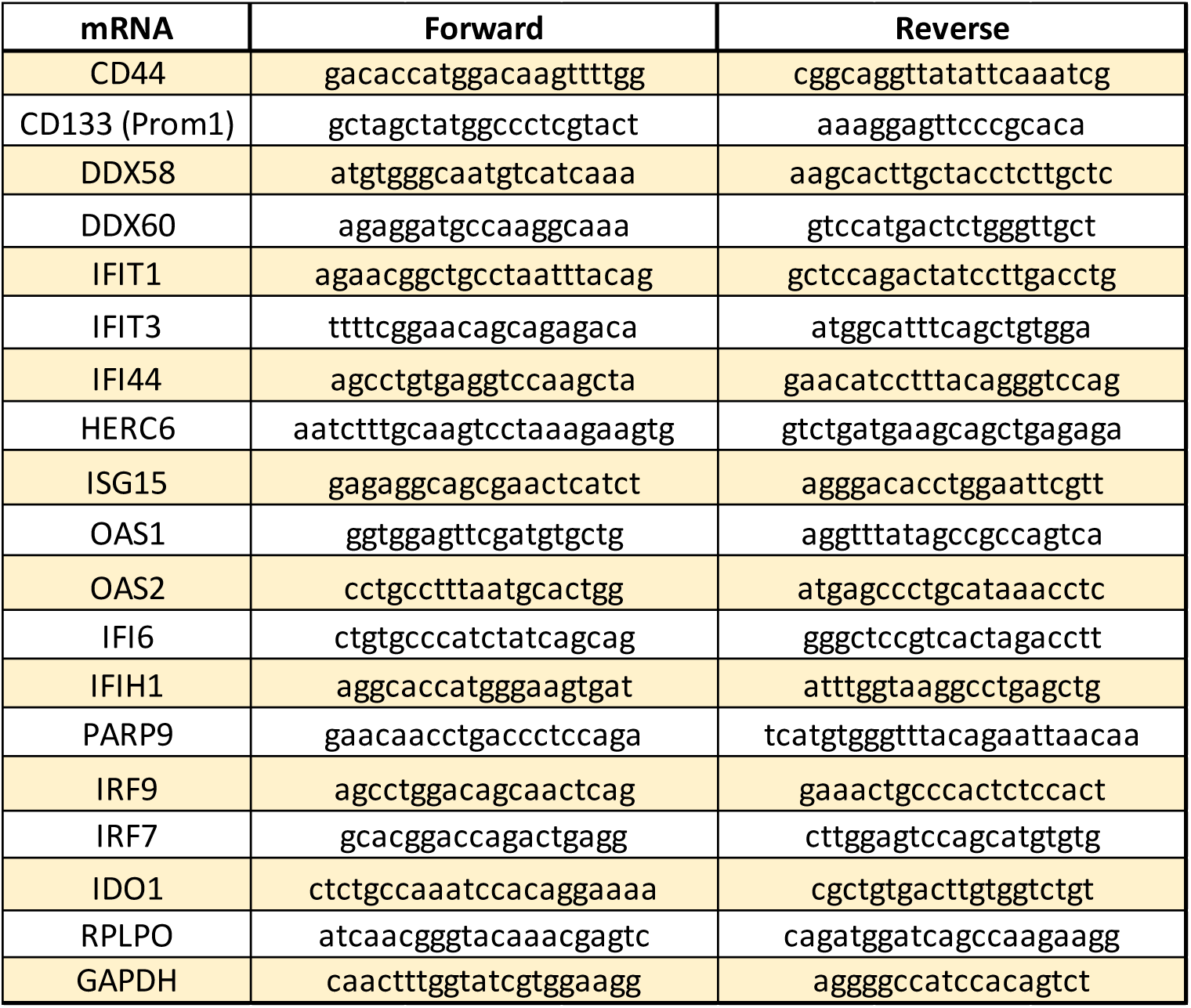
List of primer sets utilised in this study.

To assess the yield and integrity of mRNA, 2ul of each sample was taken for analysis on a Fragment Analyzer Automated CE System (Advanced Analytical) and smear analysis run using a DNF-473 Standard Sensitivity NGS Fragment Analysis Kit (Advanced Analytical). Total RNA (1ug) was processed to first-stranded cDNA using the TruSeq Stranded mRNA Library Prep Kit (Illumina), following the manufacturer’s instructions for the low-throughput protocol. First-stranded cDNA samples were processed to barcoded cDNA libraries using a Diagenode IP-Star, according to the manufacturer’s instructions for Illumina TruSeq library prep. Resulting libraries were PCR-amplified to incorporate unique indices and cleanup performed using AMPure XP Beads (Becman Coulter) according to Truseq LT protocol. Resulting libraries were quantified, pooled, and sequenced on a Nextseq 500 paired end 2×75Bp run. FASTQ files generated were de-multiplexed and adapter trimmed in the Illumina basespace and aligned to hg19 using Bowtie2 with default settings in the Basespace RNA-seq Express app. Resulting bam alignments files were exported and aligned to Refseq transcriptome annotation in Partek Genomic Suite and annotated transcripts with mapped sequence reads were quantified for relative expression in each sample and gene level RPKM (Reads Per Kilobase of transcript per Million mapped reads) counts generated for downstream analysis.

### Differential Expression Analysis

Raw count data for the IRF7 dataset was normalised using the ‘DESeq2’ R package [53] across all samples. Analysis used both IRF7 knock-down and radiation treatment individually to generate fold-change values for each gene with respect to the two main variables. These fold-change values were used as coordinates for plotting the genes (**Figure 6A**). Both IRF7 knock-down status and radiation treatment were used to generate q-values for each gene. We applied non-metric Multidimensional Scaling (isoMDS) [54] of the gene expression values to visualise similarities and differences between samples, informing data quality considerations (**Supplemental Figure 8**).

### Network Analysis

Fold change values from the normalised data generated with the DESeq2 analysis were used to assign genes into four categories: knock-down and irradiated (KD_irr), wild-type and irradiated (WT_irr), knock-down and untreated (KD_unt), and wild-type and untreated (WT_unt). A fold change threshold of 1.5 was applied to produce four non-overlapping gene-lists. These gene lists were analysed with NetNC [32] in Functional Binding Target (FBT) analysis mode, also taking HumanNet-XN network [31] as input.

The four lists were analysed separately to produce four condition-specific networks, and together to produce the ‘combined’ network. For all networks, disconnected components (clusters) with fewer than four nodes were removed.

Networks were visualised in Cytoscape (v3.7.1) [55] where functional enrichment analysis was performed using the BiNGO plugin [56]. Drug targets for coherent networks identified by NetNC were identified using the Drug-Gene Interaction database (DGIdb) [57].

Each gene was scored to quantify their network impact, according to fold change and network connectivity. Genes in the IRF7 knock-down network were scored as follows:

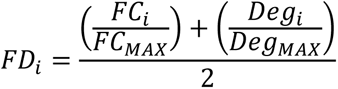

Here: FCi is the fold change value of gene i with respect to IRF7 status, FCMAX is the maximum fold change observed in the cluster, Degi is degree of gene i, and DegMAX is the maximum degree observed in the cluster.

Genes in the largest component of the IRF7 wild-type network, consisting of the viral immune response cluster, were scored as follows:

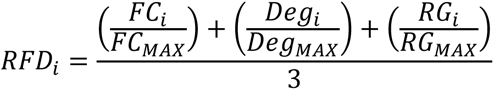

Where RGi is the number of IRF7 response genes that connect to gene i, and RGMAX is the maximum number of candidate gene connections observed in the cluster.

Ranking by FD and RFD was taken as a measure of influence over the cluster. Drug-gene interactions from DGI were followed up using the ChEMBL drug target database [58, 59]. Any genes for which the ChEMBL database has no known drug target interactions were analysed using the PubChem database [60]. If no drug target data were found in either database, the genes were assigned to group 2a. Drug target genes were analysed using ChEMBL [59] [61] SureChEMBL [56] and DrugBank (v5.0) (https://go.drugbank.com/) to assess whether the associated drug(s) satisfy Lipinski’s “Rule of Five” Lipinski [60]. Drug-target combinations that failed one or more of Lipinski’s rules were assigned to group 2b; those genes associated with drugs that passed Lipinsik’s rules were ranked by FD/RFD score. Targets with FD/RFD values>0.5 were assigned to group 3b, all others were placed in group 3a. Table 4 and Supplemental figure 12 summarise the candidate drug target groupings and the prioritisation workflow.

## Statistical Analysis

All data are shown as mean ± S.D. Tests of significance were performed by one-way ANOVA test, using Prism-GraphPad (include details). Significant changes had p-values <0.05 (* <0.05 **<0.01 ***<0.001 ****<0.0001).

## Acknowledgements

AS, MD, CAH, JMcC, LMcC, AP, FA, KB, RS, IGM, DW, SMcD, TI, NC, MDB, DM, were supported by the Belfast-Manchester Movember Centre of Excellence (MA-CE018-002), funded in partnership with Prostate Cancer UK. I.G.M. was also supported by the Norwegian Research Council (230559) and is supported by the John Black Charitable Foundation. TM and AE were supported by the Academy of Finland (grant 304667; TM), Hospital District of Helsinki, and Uusimaa (grants TYH2019235 and TYH2018214; TM) and Cancer Foundation Finland (TM). A PhD studentship from the Department for the Economy, Northern Ireland supported KMcC.

## Data availability

All data needed to evaluate the conclusions in the paper are present in the paper and/or the Supplementary Materials. The RNA-seq data has been deposited in the ArrayExpress repository (EMBL/EBI) and is accessible under accession number E-MTAB-9286.

## Author contributions

AP and FA devised and performed experiments, analysed data, and prepared the manuscript. RS, SMcD, IO and KMcC performed RNA-seq analysis. IMO and KMcC contributed to the conceptualisation, design, methodology, software, interpretation and visualization to the study of the RNA-seq data, including quality control, differential expression analysis, network analysis, prediction and ranking of candidate druggable targets. IMO supervised and provided the resources for these aspects. IMO and KMcC also undertook the data curation for the RNA-seq data and additionally contributed to the drafting and review of the manuscript. MD, MDB, TM, AE, NC, CAH, AS undertook immunohistochemistry and immune scoring of the Helsinki and Salford cohorts. AP, FA, IGM designed the study, wrote the manuscript, and analysed the data. FA, AP, JMcC, LMcC performed *in vitro* experiments. AP, FA, KB undertook the *in vivo* experiments. DM and TI contributed to the immune staining of xenograft tumours and immune micro-environment analysis. IGM supervised research, devised experiments, reviewed data, and prepared the manuscript. All authors have read and reviewed the manuscript.

## Supplementary Material

**SF1.**
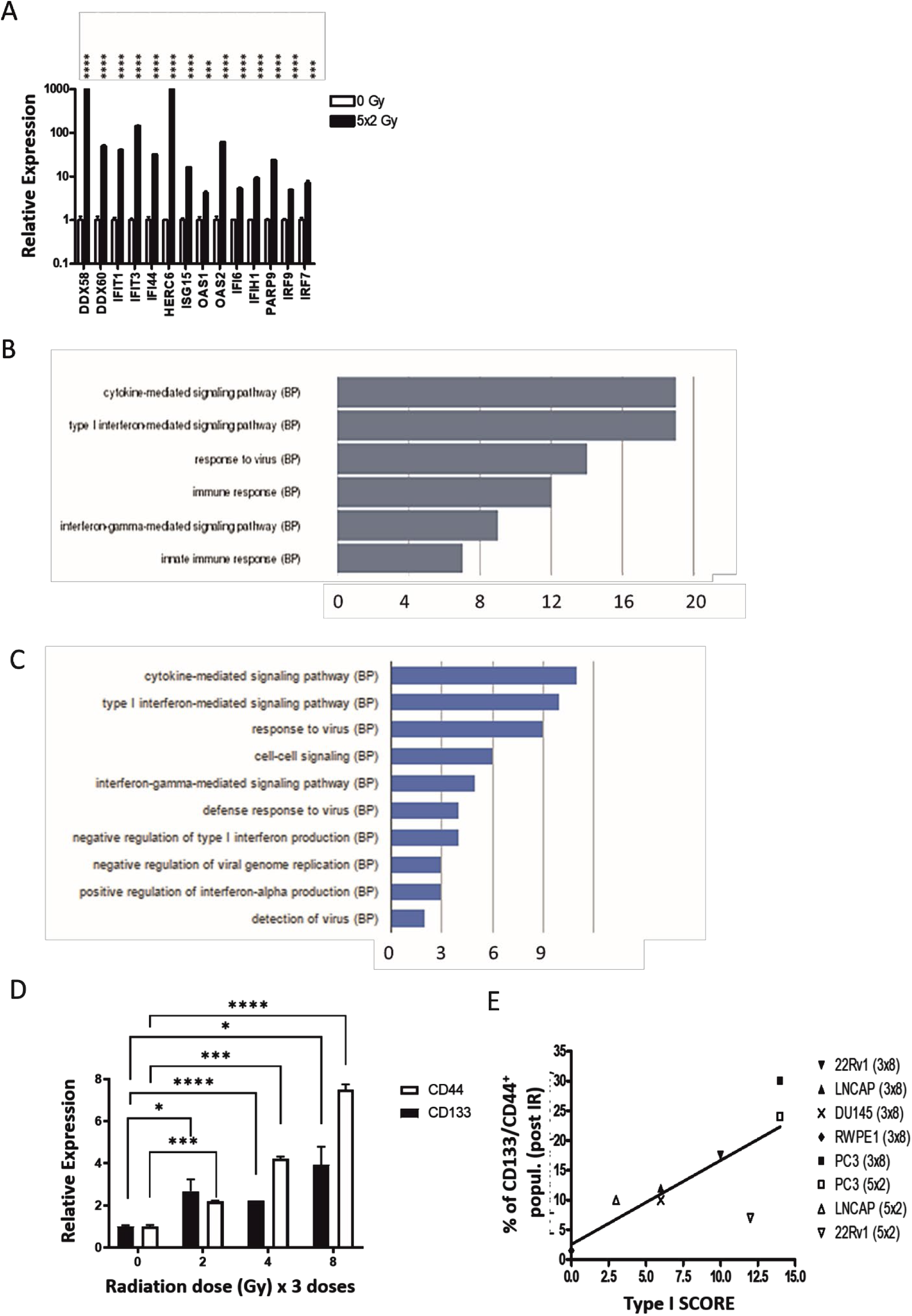
**A)** Expression of stress responsive transcripts through RT-PCR, following 5 doses of 2 Gy radiation (72 hours post final dose). **B)** Gene ontology analysis of the 81 genes identified as described in Figure 1A showed significant enrichment for cytokine mediated signalling and type 1 interferon mediated signalling. **C)** Gene ontology analysis of the 66 genes identified as described in figure 1B showed significant enrichment for cytokine mediated signalling and type 1 interferon mediated signalling. **D)** Relative expression of CD133 and CD44 with increasing doses of radiation in the PC3 cell line. **E)** Evaluation of the correlation between the expression of the stress responsive transcripts and the appearance of stem-like populations in PCa cells following irradiation. The expression of the 14 transcripts was assessed following multiple doses of radiation and normalized to un-irradiated controls. A two-fold increase in transcript expression was given a score of one up to a maximum of 14 (Type 1 score). This score was then plotted against the percentage of CD133/CD44 positive cell populations assessed by flow cytometry. A significant correlation was observed, p<0.05. **A and D-E)** All experiments represent the average of at least 3 independent experiments, p values are * <0.05 ** <0.01 *** <0.001 **** <0.0001.

**SF2.**
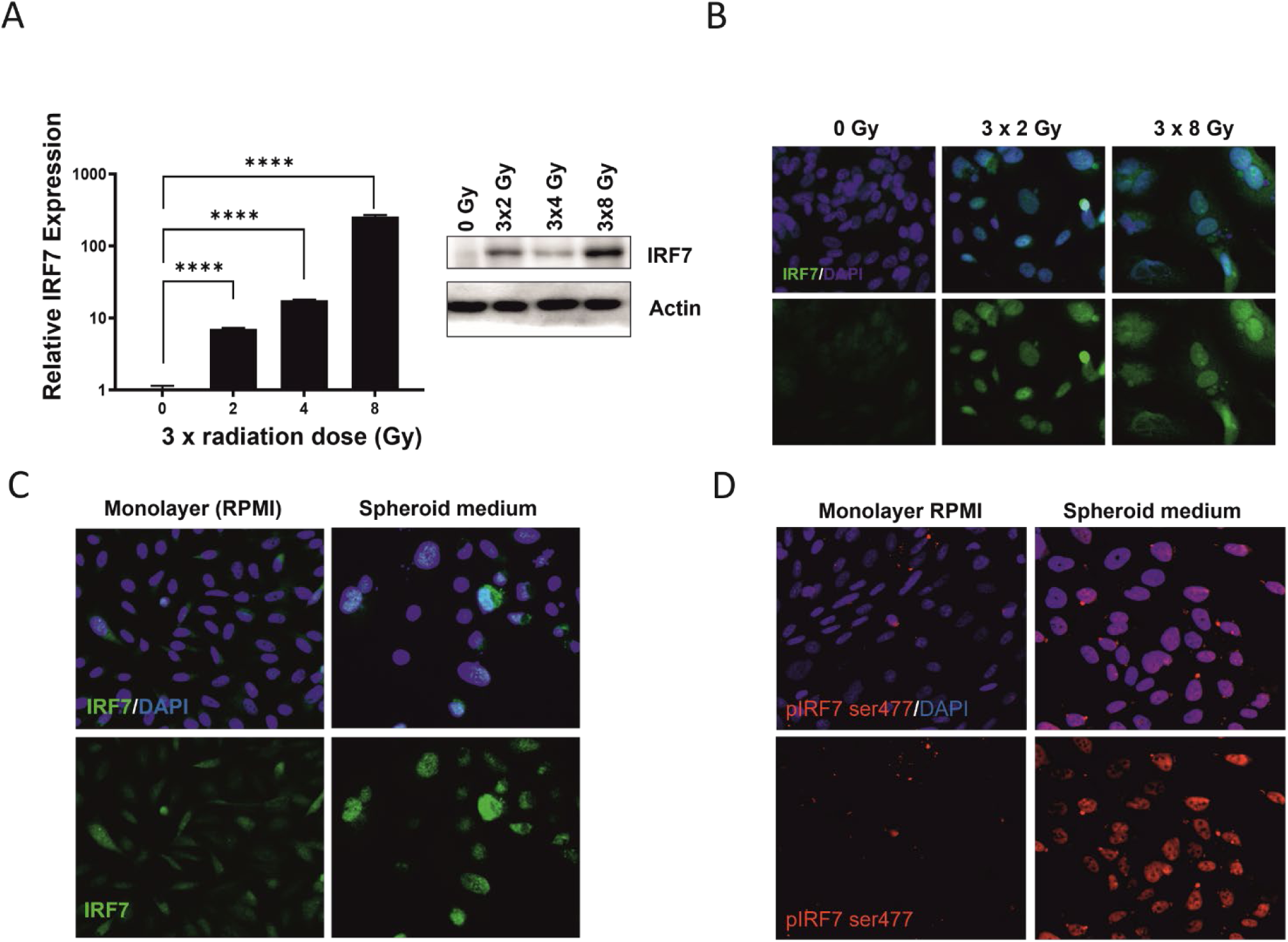
**A)** Expression of IRF7 assessed by RT- PCR and western blot following multiple doses of radiation (3 times 2, 4 or 8 Gy) at 72 hours after the last dose. The plot represents the average of n = 3 independent experiments, p values is ****<0.0001. **B)** Localization of IRF7 (green) following multiple doses of radiation (3 times 2 or 8 Gy) at 72 hours after the last dose. Nuclei are in DAPI-blue staining. **C)** IRF7 localization (green) in monolayer and spheroid forming conditions after 15 days of culture. Nuclei are in DAPI-blue staining. **D**) Detection of IRF7 phosphorylation (pIRF7 ser477, red) in cells grown under monolayer and sphere forming conditions, after 15 days of culture. Nuclei are in DAPI-blue staining. All experiments **(A-D)** are expression of n = 3 independent experiments.

**SF3.**
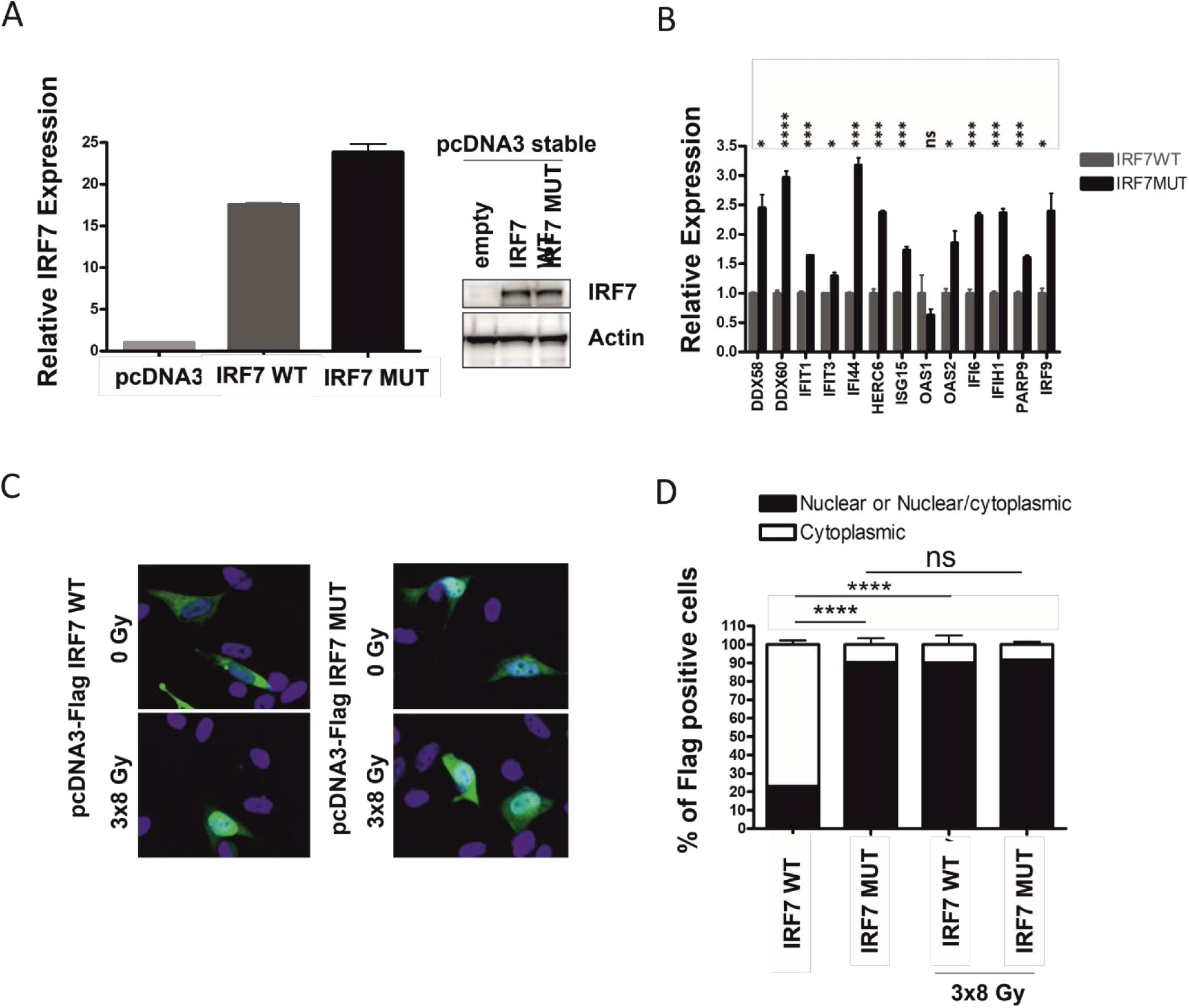
Type 1 interferon regulated gene panel is induced by phospho-mimetic mutant of IRF7 (SS477/479DD) in PC3 cells. **A)** Validation of stable overexpression of IRF7 by western blot and real- timeRT- PCR in PC3 cells. **B)** Expression of the newly identified signature through RT-PCR in mutant IRF7 expressing cells (black bars) relative to wild-type IRF7 (grey bars) in PC3 cells. **C)** Representative localisation of IRF7 (green), WT and mutant (MUT), using anti-FLAG antibodies in PC3 cells. Nuclei are in DAPI-blue staining. **D)** Quantification of nuclear and cytoplasmic localisation of IRF7 in PC3 cells, as described in (C). **(A-D)** are expression of n = 3 independent experiments, p values are * <0.05 **<0.01 ***<0.001 ****<0.0001.

**SF4.**
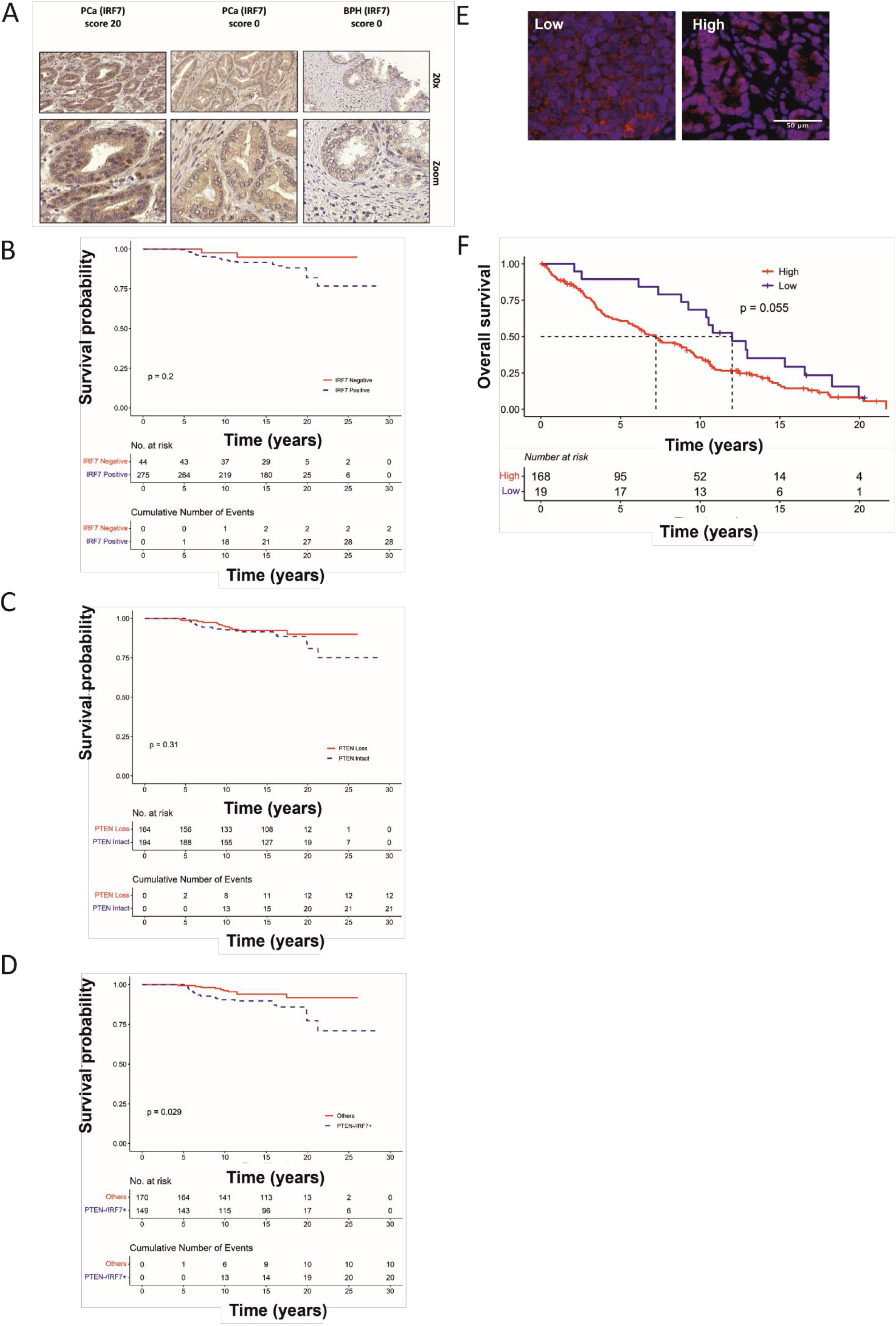
**A)** IRF7 expression in the Helsinki Prostate Cancer and TURP TMAs though immunohistochemistry staining. Representative images of scoring in benign prostatic hyperplasia (BPH) and prostate cancer (PCa) and samples collected by TURP are shown. **B)** Disease specific survival of the Helsinki TMA cohort based on nuclear IRF7 protein expression (negative vs positive), p<0.2. **C)** Disease specific survival of the Helsinki TMA cohort based on PTEN loss (overall protein expression), relative to adjacent benign prostatic epithelium (loss vs intact), p = 0.31. **D**) Cases with concomitant loss of PTEN and nuclear expression of IRF7 had a significantly poorer disease specific survival (p<0.029). **E)** Evaluation of the nuclear translocation of IRF7 in human prostate cancer tissue (red) in patients’ tumour cores from the Salford prostate cancer TURP and needle-core biopsy TMA, assessed though immunofluorescence analysis. Representative images of low and high nuclear translocation of IRF7 are shown. **F)** Patients with high Nuclear:Cytoplasmic IRF7 expression in tumour cores from the Salford prostate cancer TURP and needle-core biopsy TMA (red line) tended to have poorer overall survival compared to the remaining cohort (low, blue line). Dotted line denotes median survival. Log-rank statistic *p* values are reported (p=0.055).

**SF5.**
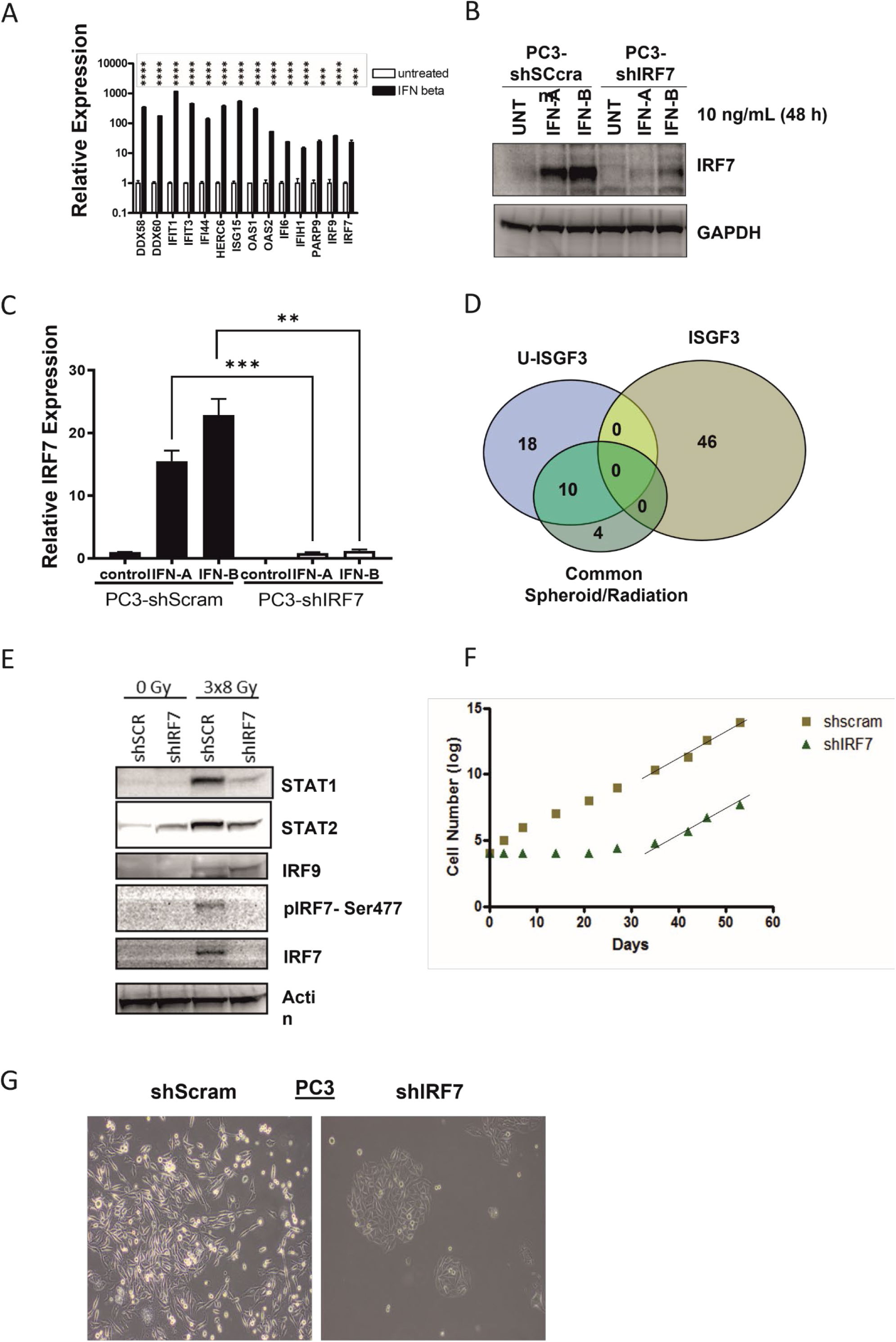
**A)** Stress responsive transcripts are interferon regulated. 48 hours interferon beta treatment of PC3 cells resulted in induction of all stress responsive transcripts as assessed through RT-PCR. The figure is expression of n = 3 independent experiments, p values are ***<0.001 ****<0.0001. **B)** Confirmation of IRF7 knockdown by western blot and **(C)** RT- PCR in PC3 cell lines. Cells were treated with Interferon alpha and beta for 48 hours to assess the degree of knockdown. **D**) Venn diagram showing the correlation of the 14 stress-responsive transcripts with those regulated by the unphosphorylated ISGF3 complex (U-ISGF3, as per literature inferred genes). **E)** Western blot analysis of IRF7 phosphorylation and status of the ISGF3 complex following radiation with 3×8 Gy. Samples were harvested 72 hours after the last dose. **F)** Growth of IRF7 knockdown PC3 cells. After an initial delay in growth, IRF7 knockdown (shIRF7) cells recover their proliferation. **G)** In the initial lag of growth IRF7 knockdowns have a cobblestone appearance compared to the fibroblastic phenotype of control cells. All experiments **(A-C and E-G)** are expression of n = 3 independent experiments, p values are **<0.01 ***<0.001.

**SF6.**
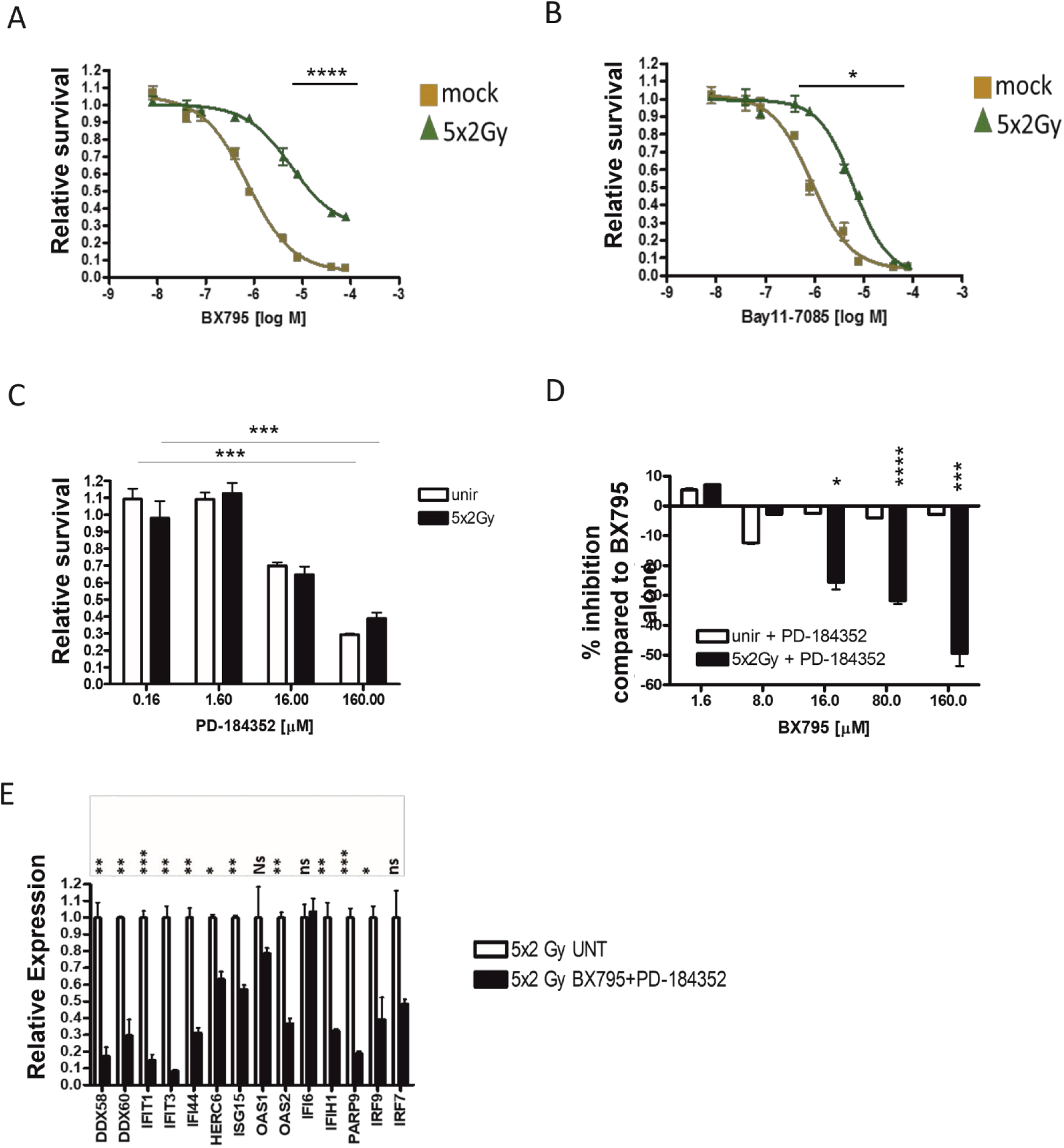
**A)** Treatment of unirradiated (mock, yellow line) PC3 cells and PC3 cells that survived 5×2Gy doses of radiation (5×2Gy, green line) with the IKKɛ/TBK1 inhibitor (BX795). Cells were treated for 72 hours, 72 hours after the administration of the last dose of irradiation. The surviving fraction is expressed as normalised on the untreated respective fraction. **B)** Treatment of unirradiated (mock, yellow line) PC3 cells and PC3 cells that survived 5×2Gy doses of radiation (5×2Gy, green line) with the IkBα inhibitor (Bay11-7085). Cells were treated for 72 hours, 72 hours after the administration of the last dose of irradiation. The surviving fraction is expressed as normalised on the untreated respective fraction. **C)** Treatment of unirradiated (unir, white bars) PC3 cells and PC3 cells that survived 5×2Gy doses of radiation (5×2Gy, black bars) with the MEK inhibitor (PD-184352). Cells were treated for 72 hours, 72 hours after the administration of the last dose of irradiation. The surviving fraction is expressed as normalised on the untreated respective fraction. **D)** Treatment of unirradiated (unir, white bars) PC3 cells and PC3 cells that survived 5×2Gy doses of radiation (5×2Gy, black bars) with the IKKɛ/TBK1 inhibitor (BX795) and the MEK inhibitor (PD-184352) in combination. Cells were treated for 72 hours, 72 hours after the administration of the last dose of irradiation. The surviving fraction is expressed as percentage of inhibition compared to the effects of BX795 alone. **A-D)** The surviving fraction after 72 hours of treatment with the inhibitors are shown. Survival was assessed by alamar blue assay. **G)** Expression through RT-PCR of stress responsive transcripts following dual treatment with BX795 and PD-184352, at 72 hours from the combination treatment. **A-G)** All experiments represent the average of at least 3 independent experiments, p values are * <0.05 **<0.01 ***<0.001 ****<0.0001.

**SF7.**
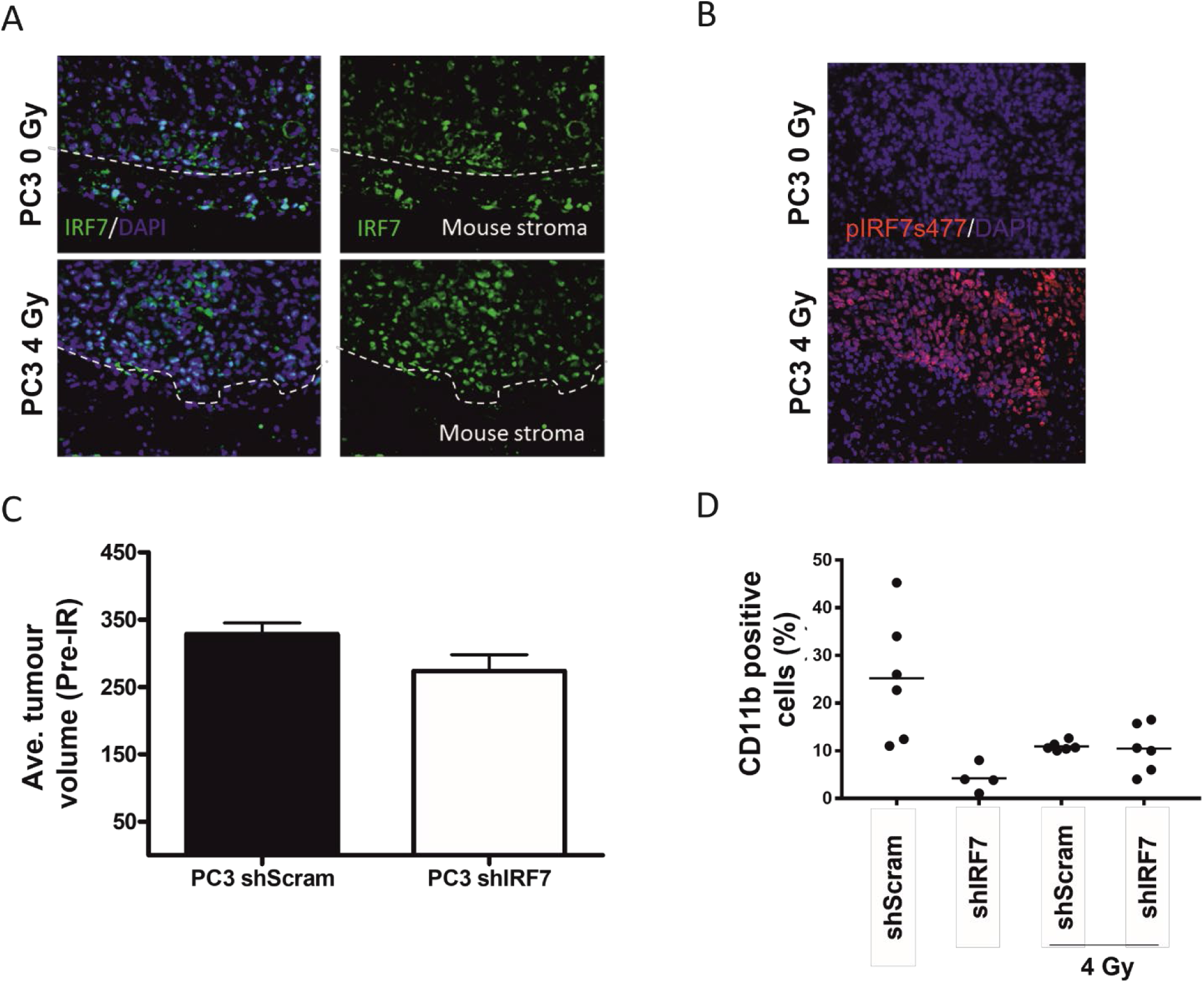
**A)** Immunofluorescent staining of IRF7 in xenografts following radiation identifies nuclear IRF7 staining. **B)** Xenografts stained with phospho-IRF7 (Serine 477) following radiation treatment. **C)** Measurements pre-irradiation of tumour volumes in control and IRF7 knockdown xenografts. **D)** Evaluation of infiltrating CD11b cells was assessed in unirradiated xenografts of control and IRF7 knockdown PC3 cells and quantified. All experiments **(A-D)** are expression of n = 3 independent experiments.

**SF8.**
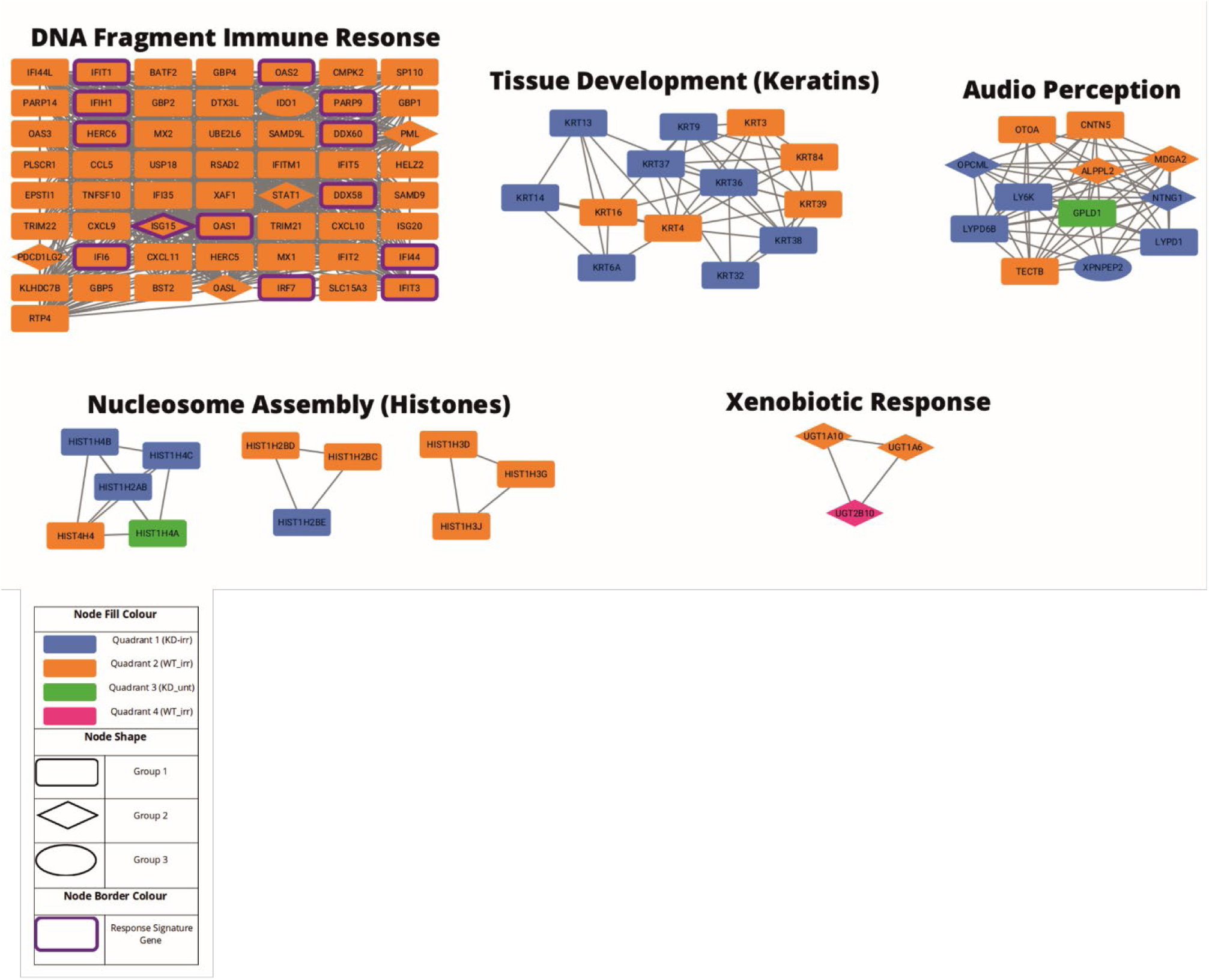
**A)** Combined network output by NetNC for genes expressed with an absolute fold change value > 1.5 in either irradiated vs untreated or IRF7 knockdown vs wild type. Node colour is determined by the condition in which the gene is present: IRF7 knock-down and irradiated (KD_irr), IRF7 wild-type and irradiated (WT_irr), IRF7 knock-down and untreated (KD_unt), and IRF7 wild-type and untreated (WT_unt). Node shape is determined by the network druggability scoring with candidate drug targets from groups 3a and 3b shown as diamonds. Clusters in the network were identified using MCL, cluster annotations are representative of functional enrichment analysis results.

**SF9.**
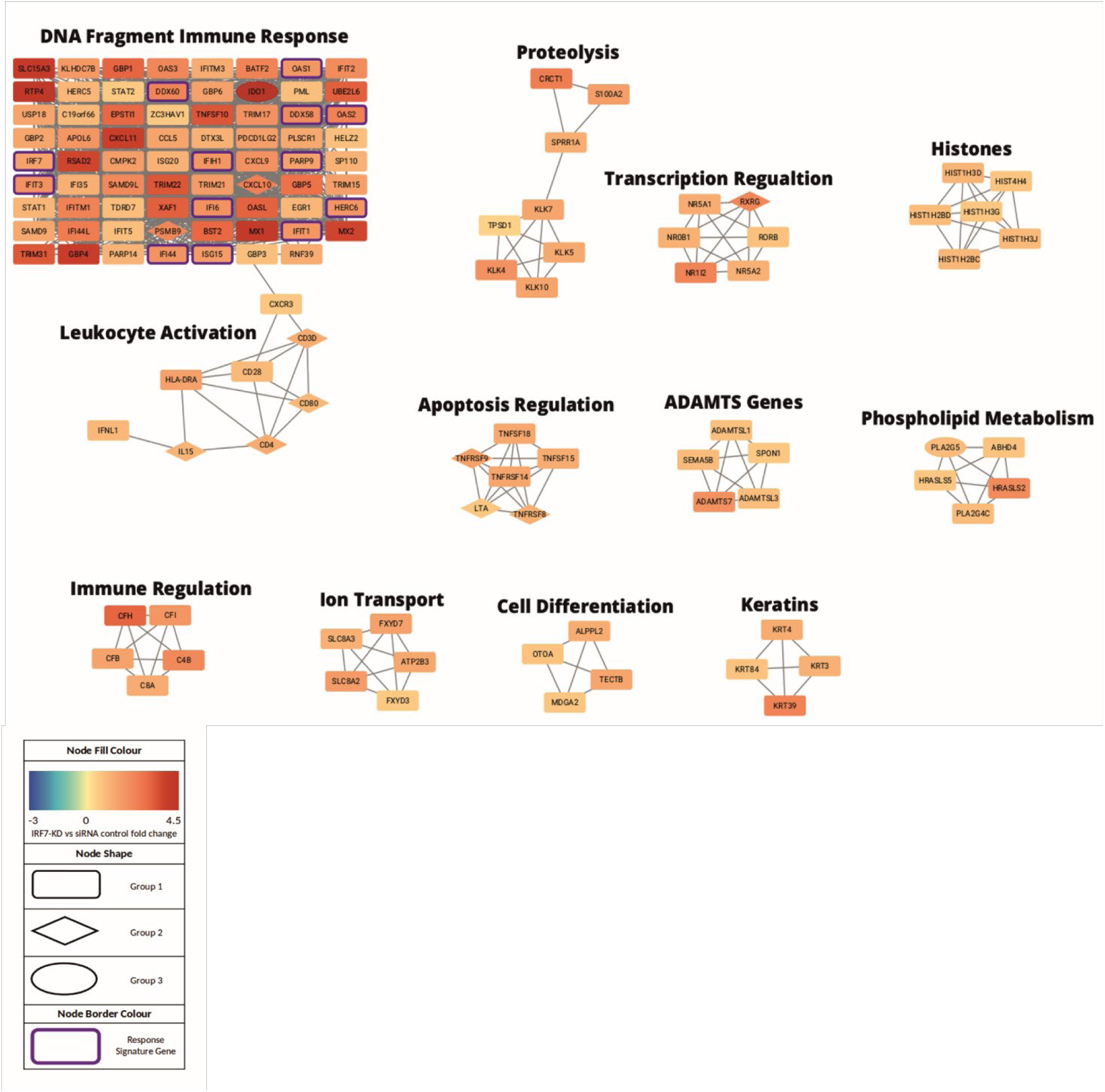
**A)** Network output by NetNC for genes at least 1.5-fold upregulated in the irradiated and IRF7 wild-type conditions. Node colour is determined by the fold-change values with respect to IRF7 status (x-coordinates in Figure 6A). Nodes with the greatest fold-change values are shown in red. Results from network drug ability analysis are indicated by node shape for candidate targets (diamonds), high-scoring candidate targets (ovals) and other genes (rectangles). Network cluster annotations are representative of significant terms from functional enrichment analysis.

**SF10.**
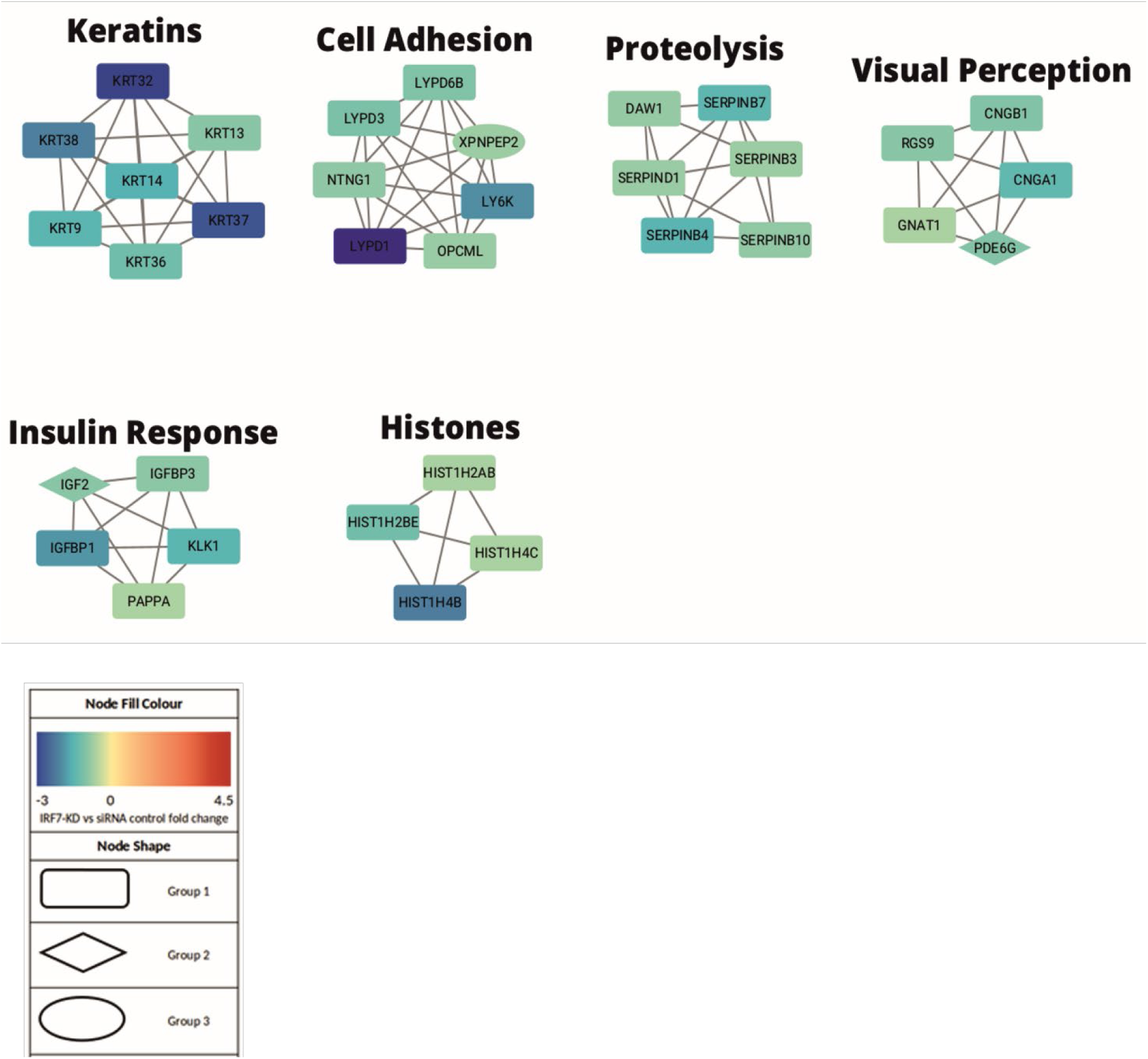
**A)** Full resulting network generated using NetNC with genes expressed in the irradiated and IRF7 knock-down with a fold change value > 1.5 for these two conditions. Node colour is determined by the fold-change values with respect to IRF7 status (x-coordinates in Figure 6A). Nodes with the greatest fold-change values are shown in blue. Results from network druggability analysis are indicated by node shape for candidate targets (diamonds), high-scoring candidate targets (ovals) and other genes (rectangles). Network cluster labels reflect significant terms from functional enrichment analysis.

**SF11.**
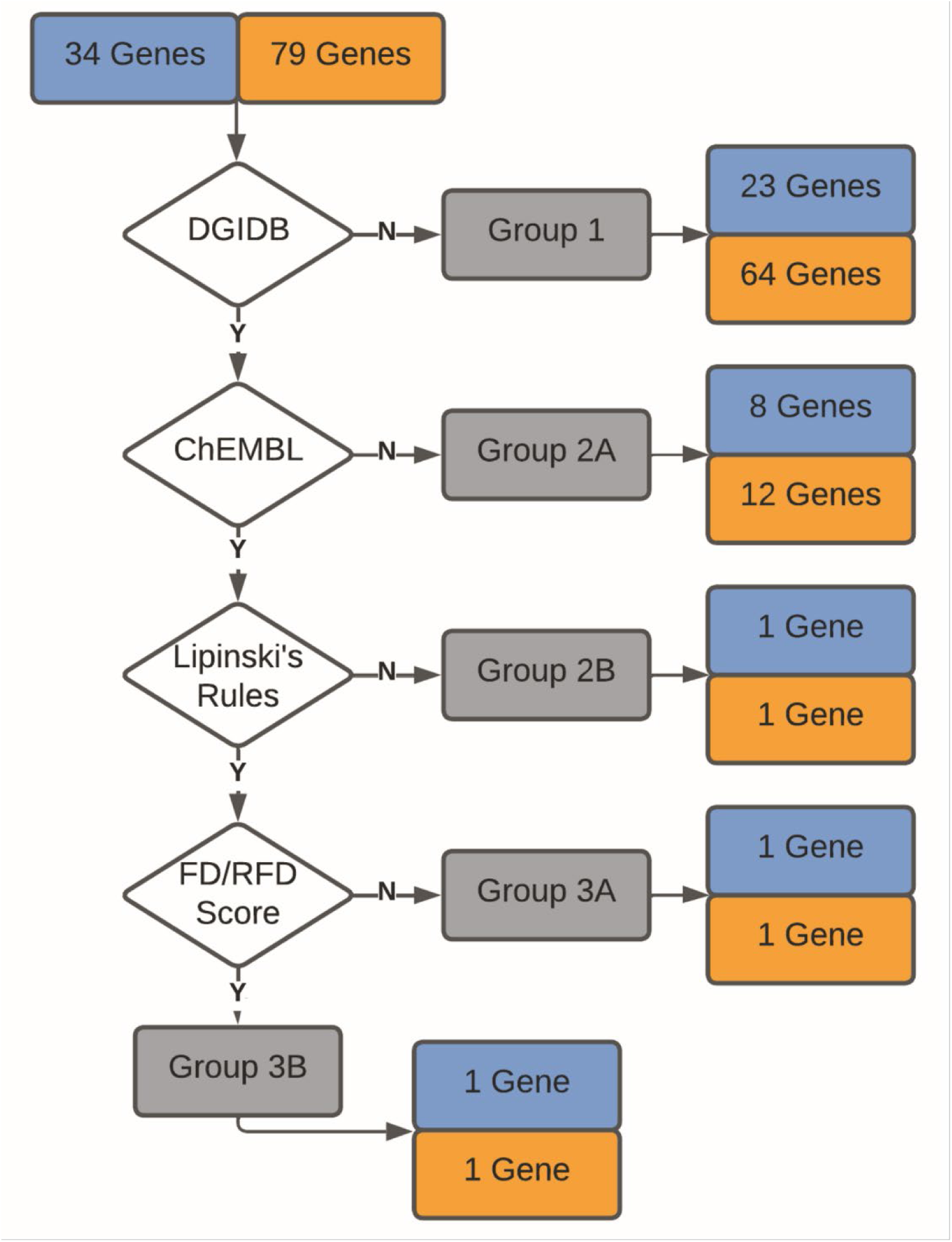
**A)** Workflow for identification and ranking of candidate target genes and associated candidate drugs from NetNC results for the irradiated IRF7 knock-down (KD_irr) and wild-type (WT_irr) conditions. NetNC returned a total of 34 and 133 genes for the irradiated KD_irr and WT_irr conditions respectively; we focussed on 79 genes in main cluster for WT_irr which contained thirteen of the IRF7-Sig genes. Associated drugs from querying DGIdb were filtered against ChEMBL and PubChem, then assessed using Lipinski’s rules. The final filter incorporated network connectivity and fold change with the FD/RFD scoring (methods), producing one high-scoring candidate target and associated drug for each network.

**SF12.**
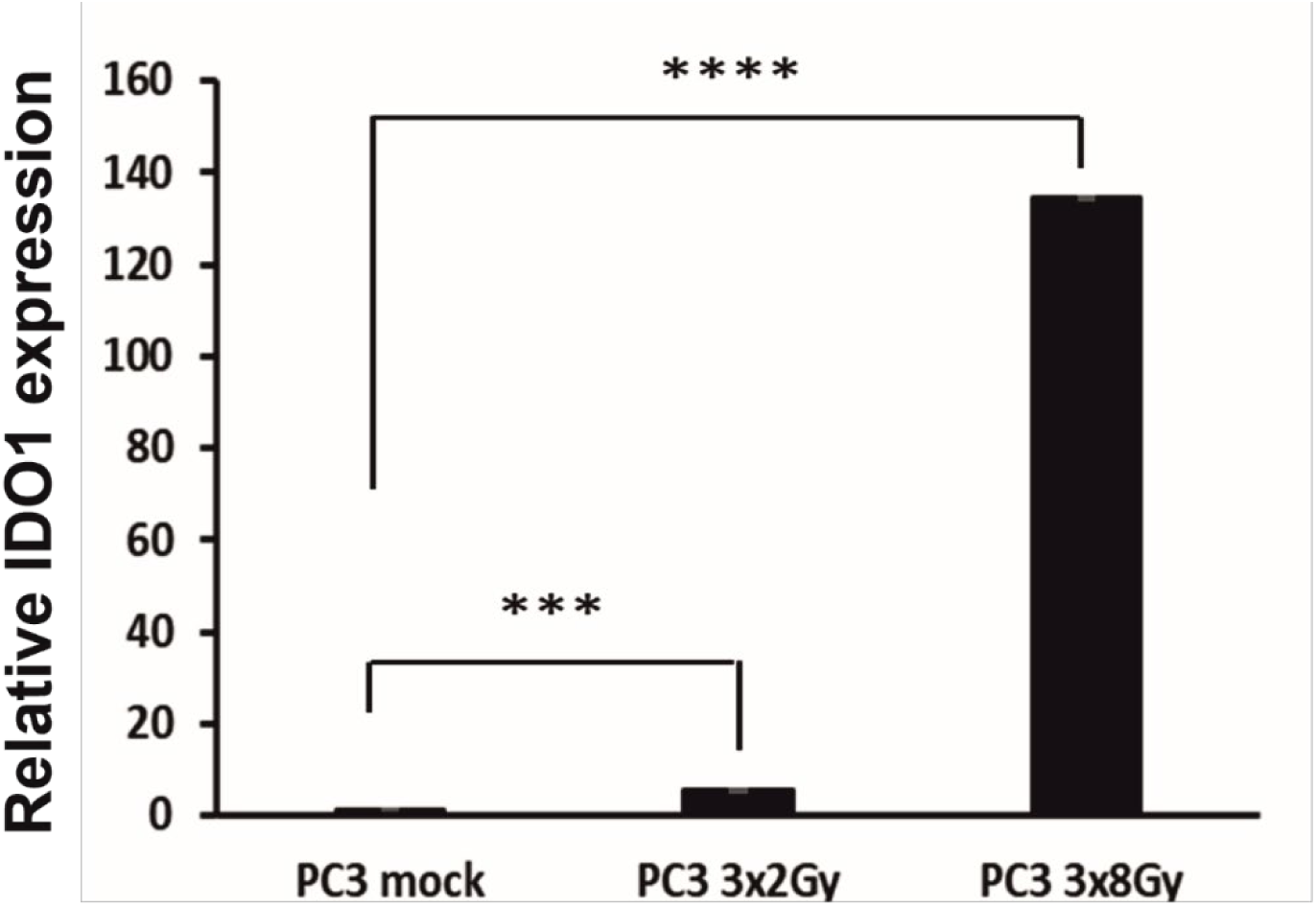
**A**) Expression of IDO1 assessed by real-time PCR following multiple doses of radiation (3×2 Gy and 3×8 Gy) compared to not irradiated cells (mock). The figure is expression of n = 3 independent experiments, p values are ***<0.001 ****<0.0001.

